# Baltic Sea microbial cohorts exhibit catabolic specialization and anabolic interdependencies across environmental gradients

**DOI:** 10.64898/2026.04.27.721049

**Authors:** Armando Pacheco-Valenciana, Felix Milke, Gerrit Wienhausen, Sarahi L. Garcia

**Affiliations:** Department of Ecology, Environment, and Plant Sciences, Science for Life Laboratory, Stockholm University, Stockholm, Sweden; Institute for Chemistry and Biology of the Marine Environment (ICBM), School of Mathematics and Science, Carl von Ossietzky Universität Oldenburg, 26129, Oldenburg, Germany; Helmholtz Institute for Functional Marine Biodiversity at the University of Oldenburg (HIFMB), Oldenburg, Germany

**Keywords:** microbial cohorts, environmental gradients, ecological traits, metabolic dependencies

## Abstract

Microbial communities are structured by environmental gradients and metabolic interactions, yet the genomic characteristics and metabolic functions of co-occurring populations remain underexplored. Here, we investigated co-occurring microbial cohorts across the Baltic Sea, a system characterized by strong salinity, temperature, and oxygen gradients. For this, we used a genomic catalog consisting of 701 species-representative genomes to recruit reads from 112 metagenomes and infer cohort structure, environmental distributions, and metabolic potential. We identified nine microbial cohorts that showed strong associations with environmental gradients, indicating deterministic assembly. Cohorts differed markedly in genomic traits, with the most abundant and prevalent taxa associated with smaller, streamlined genomes, while a low-oxygen cohort with larger genomes contributed disproportionately to nitrogen and sulfur transformations. Across cohorts, biosynthetic potential was unevenly distributed. Amino acid biosynthesis pathways were frequently complete, whereas B-vitamin pathways were typically incomplete and rarely encoded in full by individual genomes. Metabolites with low pathway completeness showed consistent taxonomic partitioning, with biosynthetic capabilities distributed across taxa rather than collectively encoded within cohorts. Together, these results show that Baltic Sea microbial cohorts are ecologically structured assemblages whose genomic repertoires reflect catabolic specialization and anabolic interdependencies. Our findings highlight microbial cohorts as a useful framework for linking environmental gradients, genome traits, and the organization of metabolic functions in natural microbial communities.

## INTRODUCTION

Five quintillion (5x10^30^) of prokaryotes are the engines that drive Earth’s biogeochemical cycles, keeping the planet habitable as we know it (Falkowski et al., 2008; Whitman et al., 1998). While estimates of their diversity surpass the millions, a global census shows that we have sequenced only a small fraction of them (Parks et al., 2026; Schloss et al., 2016). Even fewer microbes are available in culture and have been studied in depth (Lewis et al., 2021; D. Wu et al., 2025). While studying microorganisms in pure culture leads to a deep understanding of their physiology and mechanistic molecular strategies for growth and survival, there are emergent properties of life that can only be studied when microorganisms are surrounded by one another. As such, microorganisms can be studied at different levels of organization (Gralka, 2023; Milke et al., 2026), such as in their natural populations (Freel et al., 2025), in co-cultures (Wienhausen et al., 2024), functional consortia (Pacheco-Valenciana, Tausch, et al., 2025), or communities in nature (Giordano et al., 2024). To accomplish the study of some of these levels, approaches such as genome-resolved studies have become fundamental in microbiology.

Advances in DNA sequencing over the past few decades have facilitated the reconstruction of microbial genomes and have enabled a more direct investigation of microbial structure and metabolic potential from environmental samples (Albertsen et al., 2013; Eren & Banfield, 2024; Tyson et al., 2004). One approach to infer community structure is through co-occurrence network analysis (Friedman & Alm, 2012). Such networks have been found to exhibit modular structures (Ma et al., 2020; Milke et al., 2023; Newman, 2006), where groups of microorganisms co-occur across environmental gradients. These microbial groups are currently referred to by various names, including “modules”, “metabolically cohesive consortia”, “functional cohorts”, and “guilds” (Milke et al., 2023; Mondav et al., 2020; Pascual-García et al., 2020; G. Wu et al., 2021), but here we will refer to them as cohorts. Despite the terminological differences, their modularity, compositional stability, distinct ecological preferences, and ubiquity suggest that cohorts are a meaningful level of microbiological organization (Milke et al., 2026), composed of co-occurring species whose associations are shaped by joint environmental preferences and potential biotic interactions.

At the same time, genome-resolved metagenomic studies have advanced our understanding of microbial functions (Castelle et al., 2018; Wrighton et al., 2012). In aquatic environments, biogeochemical cycles can be sustained through metabolic handoffs, with thousands of species contributing only subsets of sequential redox transformations rather than encoding entire pathways individually (Anantharaman et al., 2016). In fact, many abundant aquatic microorganisms exhibit small genome sizes together with a reduced biosynthetic repertoire (Giovannoni et al., 2005; Rodríguez-Gijón et al., 2025; Swan et al., 2013). This phenomenon, known as genome streamlining, occurs across multiple marine, brackish, and freshwater lineages and has been further associated with increased anabolic interdependence (Giovannoni et al., 2014; Pacheco-Valenciana, Tausch, et al., 2025; Rodríguez-Gijón et al., 2025). The ecological implications of genome streamlining are partly explained by the Black Queen hypothesis, which posits that microorganisms lose costly metabolic functions as long as the respective metabolite is synthesized by other organisms and available in the environment (Morris et al., 2012). Within this framework, microbial populations with reduced metabolic capabilities may benefit by depending on shared metabolites generated by the community (Giordano et al., 2024; Kost et al., 2023).

Biosynthesis of essential metabolites is particularly important in this context. Several genomic studies show that many microorganisms lack biosynthetic pathways for compounds such as amino acids, vitamins, or cofactors (Gómez-Consarnau et al., 2018; Kim et al., 2021; Ramoneda et al., 2023). Although amino acid biosynthesis pathways are more often retained in many microbial genomes, B-vitamin biosynthesis pathways are often incomplete or absent (Rodríguez-Gijón et al., 2025). B-vitamins act as enzymatic cofactors, resulting in a low cellular demand, which may contribute to the widespread reliance on environmental sources rather than maintaining full biosynthetic pathways (Sañudo-Wilhelmy et al., 2014). Moreover, biosynthetic potential is not strictly binary. Some microorganisms can complete pathways if some intermediates are available, while others rely on the uptake of the final metabolite (Hong et al., 2025; Klier & Anantharaman, 2026; Wienhausen et al., 2024). As a result, biosynthetic capabilities in microbial communities probably exist along a spectrum, ranging from more to less independent (Pacheco-Valenciana, Tausch, et al., 2025). However, whether biosynthetic pathway steps are partitioned across genomes within ecologically defined microbial cohorts remains unresolved.

In this study, we investigate the distribution of key genes involved in biogeochemical cycles as well as biosynthetic pathways across microbial cohorts in the Baltic Sea. The Baltic Sea spans large environmental gradients, foremost salinity and oxygen, and is therefore ideal to investigate cohort distribution and function. For this, we mapped a Baltic Sea catalog of 701 species-representative genomes to 112 Baltic Sea metagenomes from different locations to infer genome abundances and co-occurrence-based cohort associations. We test how microbial cohorts assemble along environmental gradients and whether metabolic functions and biosynthetic capabilities across genomes express cohort-specific patterns. While genome-resolved metagenomics has inherent limitations, including incomplete annotations, biases in gene assignment, and incomplete genomes reconstructed from metagenomes, our main conclusions were robust despite genome incompleteness. The goal is to observe trends that could inform us about functional diversity and complementarities between and within cohorts. Overall, we show that individual cohorts carry a functional signature that aligns with their abiotic preferences. Moreover, interdependencies for three amino acids and three B-vitamins seem to be widespread across members of all cohorts. Our findings open the discussion of emergent functional diversity of microorganisms when living surrounded by their co-occurring microbes, emphasizing the ecological importance of interdependence.

## MATERIALS & METHODS

### Metagenomic dataset and genome catalog

We used a published metagenome-assembled-genome (MAG) catalog from the Baltic Sea that includes 1603 genomes (medium to high quality) that were dereplicated at 95% average nucleotide identity (ANI), resulting in 701 species-clusters (Pacheco-Valenciana, Tausch, et al., 2025). For each of these species-clusters a genome representative was chosen with the best quality. Each genome representative is used as a proxy for a species. These genomes were used to recruit reads from 112 metagenomic samples (**Figure 1A**) across the Baltic Sea, spanning four independent studies, with multiple geographic locations and water depths (Alneberg et al., 2018, 2020; Larsson et al., 2014; Pacheco-Valenciana, Tausch, et al., 2025). Metadata for the 701 species-representative genomes, including taxonomic classification, genome quality metrics, and cohort assignment, are provided (**Supplementary Table S1**), together with metadata for the 112 metagenomic samples, including accession number, study origin, and environmental variables (**Supplementary Table S2**).

**Figure 1.**
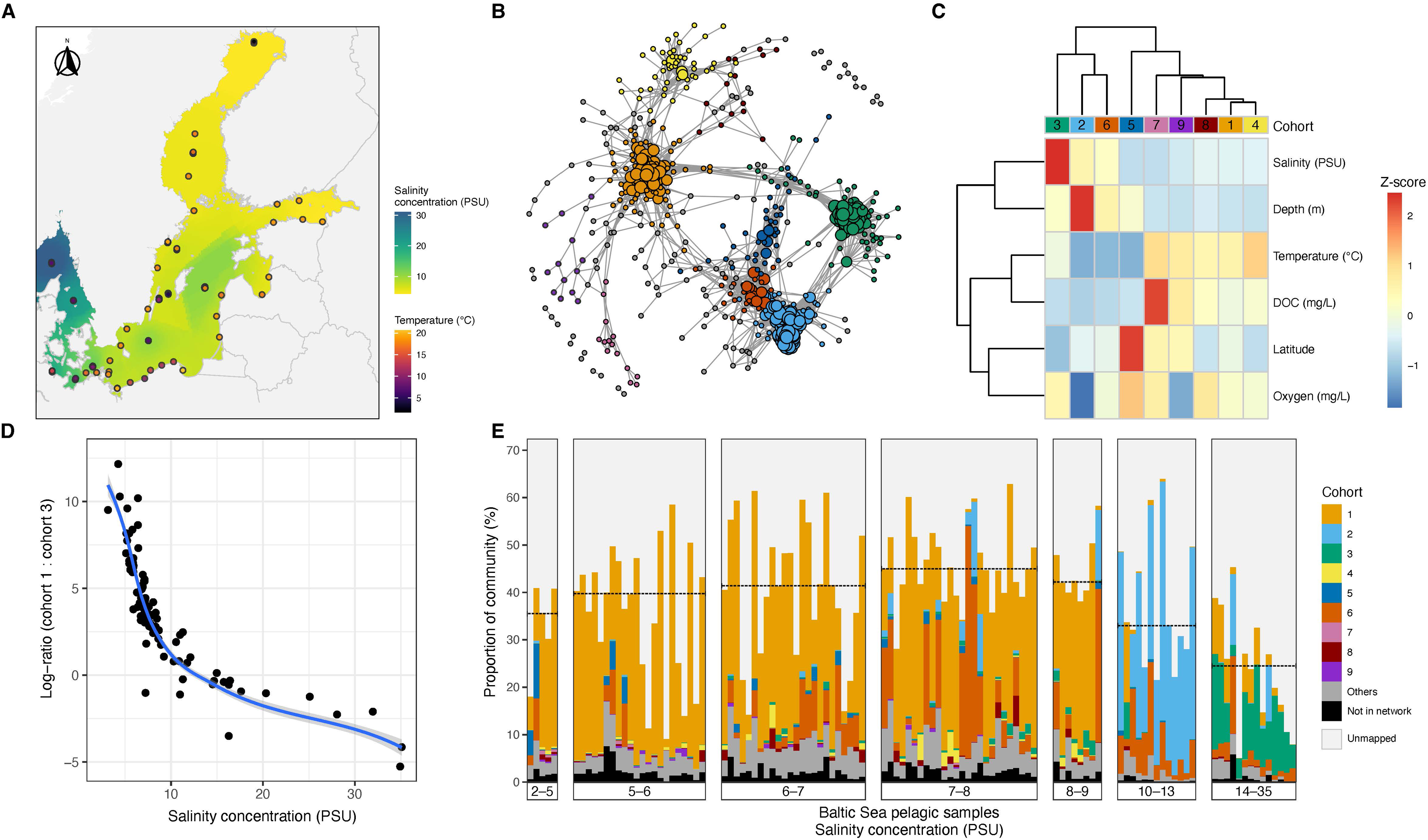
Environmental structuring and cohort-level organization of Baltic Sea microbial species. (A) Map of the Baltic Sea showing the location of publicly available pelagic metagenomic samples (n = 112). Points indicate sampling sites. The color of the water represents salinity concentration (PSU), and the point color denotes in situ temperature (°C). (B) Species co-occurrence network inferred from relative abundance correlations across all environmental samples (593 species). Each node represents a species, and edges indicate the connection among them. Nodes are colored by cohort. (C) Heatmap shows environmental preferences of cohorts, calculated as z-scored average environmental conditions weighted by relative abundances of each cohort. Columns are clustered by similarity. (D) Log-ratio of relative abundance between Cohort 1 and Cohort 3 across the salinity gradient. Each point represents one metagenome, and the fitted loess-curve illustrates the continuous shift in dominance along increasing salinity. (E) Stacked bar plots showing the relative contribution of microbial cohorts to total community composition in each metagenome (n = 112), ordered by increasing salinity (PSU). Colors represent the cohorts identified in the network. “Others” denotes genomes not assigned to any cohort, “Not in network” indicates genomes excluded from network analysis, and “Unmapped” represents reads not mapping to any of the 701 reference species collection. Dashed lines indicate the median proportion of mapped reads within each salinity category.

### Estimation of species relative abundance and prevalence

Briefly, the relative abundances were estimated by read recruitment using Strobealign (v0.14.0) (Sahlin, 2022). The resulting BAM files were processed within the Anvi’o platform (v7.1) (Eren et al., 2015). BAM files were first prepared (sorted) using the program ‘anvi-init-bam’, and genome coverage across samples was then calculated with ‘anvi-profile-blitz’. For each sample, the relative abundance of each species was calculated by dividing the genome’s mean coverage of the inner quartiles (q2q3_cov) by the sum of the mean coverage (of the inner quartiles) of all genomes in that sample. Relative abundance values for all species across samples are provided (**Supplementary Table S3**). Prevalence was defined as the proportion of metagenomic samples in which a species was detected. A species was considered present in a sample when its estimated relative abundance was greater than zero. Mean relative abundance and prevalence values calculated across all samples are reported for each species (**Supplementary Table S1**).

### Co-occurrence network construction

We inferred a co-occurrence network from relative abundance data of MAGs using the SParCC algorithm, implemented via the computationally efficient FastSpar adaptation. SparCC infers correlation metrics from compositional data, reflecting both potential biotic interaction between MAGs and shared environmental preferences. For downstream analyses, we focused only on positive correlations, since we are interested in groups of MAGs that share the same spatio-temporal distribution, here defined as microbial cohorts. To account for false discovery rates, we bootstrapped network inference and recalculated correlation metrics based on shuffled datasets. The resulting *p*-values are defined as the proportion of shuffled datasets whose correlation is equal or higher than the observed value, as recommended by the original SparCC publication (Friedman & Alm, 2012). We discarded all correlations with r-values < 0.6 and *p*-values > 0.05. The resulting network was further analyzed in R using the igraph package. Clusters were computed based on edge betweenness (Newman & Girvan, 2004), and all clusters with equal or less than six members were discarded to remove spurious or insufficiently sampled clusters. The resulting clusters are henceforth termed cohorts. Modularity was calculated from the resulting cluster memberships. We computed the probability to which clusters deviate from random expectations through bootstrapping (n = 1000) by rewiring the observed network using 10 times the number of vertices, as done elsewhere (Milke et al. 2026). The probability of being different from random expectations was calculated as the proportion of bootstrapped iterations that produced equal or higher modularity than observed. For MAGs assigned to cohorts, we calculated their degree of connectedness (node degree), defined as the number of edges connected to each MAG in the inferred co-occurrence network.

### Detection of carbon, nitrogen, and sulfur metabolic genes

Genes involved in carbon, nitrogen, and sulfur transformations were identified across all species-representative genomes using METABOLIC (v4.0) (Zhou et al., 2022). This pipeline integrated Hidden Markov Model (HMM) searches against curated metabolic marker gene databases, including KEGG ortholog profiles and carbohydrate-active enzyme annotation from the dbCAN database (Zhang et al., 2018). Each genome was analyzed using the genome-scale workflow (METABOLIC-G), which predicts metabolic functions based on HMM profile searches and curated pathway rules. From the full METABOLIC output, we extracted a subset of genes associated with key transformations in carbon, nitrogen, and sulfur cycling. Presence or absence of these genes was compiled into a table for downstream analysis (**Supplementary Table S4**).

### KEGG module completeness analysis for amino acids and vitamins

Functional annotations for KEGG Orthologs (KOs) and estimates of metabolic potential were obtained using the Anvi’o platform (v7.1) (Eren et al., 2015). Each genome was annotated against the KEGG KOfam database (Aramaki et al., 2020), and metabolic capabilities were inferred using the ‘anvi-estimate-metabolism’ program (Veseli, Chen, et al., 2025). To assess biosynthetic potential for amino acids and vitamins, a curated set of custom KEGG-based modules representing biosynthetic pathways for these metabolites was implemented (Pacheco-Valenciana, Tausch, et al., 2025). These modules were defined by selecting KEGG Orthologs directly from KEGG pathway maps and encoding logical relationships between pathway steps (AND/OR rules) to represent alternative biosynthetic routes. These curated modules were designed to reflect *de novo* biosynthetic potential under the specific pathway definitions used here, rather than all possible salvage, uptake, or alternative downstream routes that could also contribute to metabolite production. Where multiple KEGG modules produced the same metabolite, they were consolidated into a single custom module. For vitamin B_12_, the completeness of the custom module was evaluated separately for the aerobic and anaerobic biosynthetic routes, each of which combined corrin ring biosynthesis with nucleotide loop assembly into a single route-specific custom score. In analyses requiring a single summary value per genome, B_12_ completeness was defined as the best-route completeness, calculated as the higher of the aerobic and anaerobic custom module values. Custom module definitions are publicly available in the associated repository (Veseli, Pacheco-Valenciana, et al., 2025).

Because genome incompleteness affects the inference of pathway absence and presence differently, we applied different genome filters depending on the question being asked. Analyses of fully absent pathways (0% completeness) were restricted to high-quality genomes (>90% completeness and <5% contamination), because missing genomic regions in lower-quality assemblies can artificially inflate the number of pathways inferred as absent. In contrast, analyses of fully complete pathways (100% completeness) included all genomes, since detection of the full set of required genes provides positive evidence for biosynthetic potential even in a partially recovered genome. Thus, heatmaps summarizing the proportion of genomes with no detected pathway steps were calculated using only high-quality genomes, whereas heatmaps summarizing the proportion of genomes encoding complete pathways were calculated using all genomes.

For the present study, we analyzed module completeness values ranging from 0 to 1 for the curated set of amino acid and B-vitamin biosynthesis modules. Rather than treating biosynthetic capacity as a strictly binary trait, we interpreted pathway completeness as a gradient, consistent with previous analyses showing that variation in biosynthetic completeness explains major differences among Baltic Sea species (Pacheco-Valenciana, Tausch, et al., 2025). For visualization, genomes were assigned to five biosynthetic completeness categories: 0%, 1–35%, 36–66%, 67–99%, and 100%. These categories were defined to provide an interpretable summary of pathway completeness values across genomes, while distinguishing complete absence (0%) from complete pathways (100%). Importantly, genomes in the 0% category were interpreted as lacking all detected steps for a given curated module under our annotation framework, rather than being directly classified as auxotrophs. Additionally, we analyzed gene-level presence–absence data for individual steps within the KEGG modules M00924 (anaerobic corrin ring biosynthesis), M00925 (aerobic corrin ring biosynthesis), and M00122 (nucleotide loop assembly, NLA), which together represent the anaerobic and aerobic routes of the B_12_ biosynthesis pathway. Completeness values for all genomes and custom pathway modules are provided (**Supplementary Table S5**), and the presence–absence matrix of individual pathway genes for B_12_ is included (**Supplementary Table S6**).

### Statistical analyses

We used R (v4.4.0) (R Core Team, 2024) and RStudio (RStudio Team, 2024) to perform all statistical analyses. To assess differences among more than two groups, we used the Kruskal–Wallis rank-sum test (Kruskal & Wallis, 1952). When the Kruskal–Wallis test showed significant differences (*p* < 0.05), we performed pairwise comparisons with Dunn’s test and applied the Bonferroni correction for multiple testing (Dunn, 1964). Spearman’s rank correlation coefficient (ρ) was used to assess associations between variables (Spearman, 1987), and the corresponding *p*-values are reported for all correlations.

We fitted generalized additive models (GAMs) (Hastie & Tibshirani, 1986) to infer how well cohort abundance ratios can be predicted from the three environmental parameters, salinity, temperature, and oxygen concentration. For this, we assumed Gaussian errors and used an identity link function. Smooth functions were fitted using thin plate regression splines, and smoothing parameters were selected by generalized cross-validation. The total estimated degrees of freedom varied between 4.76 and 14.71. All analyses were conducted in R using the mgcv package (Wood, 2011).

## RESULTS

### Microbial cohorts in the Baltic Sea assembled along environmental gradients

To identify groups of species that consistently co-occur across the Baltic Sea, we constructed a co-occurrence network using the relative abundance patterns of all 701 species across samples. The co-occurrence network analysis retained 85.7% of the species (n = 593) and revealed nine microbial cohorts, with an average modularity of 0.707 (**Figure 1B**). The cohort sizes varied significantly. Three large cohorts (Cohorts 1 to 3) included between 94 and 115 species. Three medium cohorts (Cohorts 4 to 6) had 35 to 51 species, and three small cohorts (Cohorts 7 to 9) had just 13 to 14 species. Of the 593 species in the network, 113 nodes clustered into very small groups, representing ≤ 5 nodes, and were labeled as “Others”. In total, 108 species did not express positive co-occurrences and were therefore not included in the network.

To assess how well the inferred cohorts represented the entire metagenomic communities, we then quantified read mapping across samples. On average, 38.63% of metagenomic reads per sample mapped to the BalticMAG catalog, with the highest representation in intermediate-salinity regions (7–8 PSU) and the lowest in high-salinity regions (14–35 PSU; **Figure 1E**). On median, 45% of reads mapped to the MAG catalog at salinity 7–8 PSU. When considering the full metagenomes, the defined cohorts accounted for 15.44% to 34.92% of total reads across all samples (mean = 30.16%). Relative only to reads mapping to the species catalog, cohorts collectively represented an average of 83.9% of the mapped community.

The microbial cohorts showed clear environmental preferences (**Figure 1C**). For example, Cohort 1 was highly abundant in low-salinity samples (2–9 PSU), representing on average approximately 24.9–32% of all reads in those samples. Cohort 2 peaked at intermediate salinities (10–13 PSU) and had a strong preference for lower oxygen and greater depths. In contrast, Cohort 3 was predominantly found in samples with high salinity concentrations (14–35 PSU), representing, on average, 10.2% of the reads. Cohort 4 preferred higher temperatures, and Cohort 5 preferred high latitude and low salinity. Cohort 6 became more abundant at salinities above 6 PSU and preferred greater depths and colder temperatures (**Figure 1E**).

Salinity clearly affected the composition of microbial cohorts, leading to a continuous change between the three largest cohorts (Cohorts 1, 2, and 3). This gradual change led to a non-linear trend between sample-wise log-ratios of Cohorts 1 & 3 and environmental conditions (**Figure 1D**). Cohort 1 was more common at lower salinities, while Cohort 3 became more dominant at higher salinities **(Figure 1D**). Focusing on log-ratios is particularly interesting, as it avoids compositionality biases of sequencing data. This led us to further investigate the predictive performance of cohort composition based on the three most important environmental gradients: salinity, temperature, and oxygen. Accounting for non-linear relationships using generalized additive models (GAMs), we found highly predictive models for a set of ratios covering all 9 microbial cohorts (deviance explained 76.8–94.5%; **Supplementary Table S7**). This strong predictability of microbial cohort composition, based solely on salinity, temperature, and oxygen, underlined the deterministic assembly of microbial cohorts and likely reflects niche specialization across cohorts.

### Baltic Sea cohorts with smallest genome size are also most abundant and prevalent

To understand how this niche specialization translates into genomic organization at the species level, we focused on the six largest cohorts (Cohorts 1–6), each comprising more than 30 species and also accounting for most of the relative abundance across samples (**Figure 1E** and **Figure 2B**). Smaller cohorts (Cohorts 7–9), with fewer than 15 species, were not included in the subsequent comparative analysis due to the limited number of genomes available for robust comparisons. Although the investigated cohorts were predominantly composed of high-quality genomes, genome completeness differed significantly across cohorts (Kruskal–Wallis, *p* = 0.005), with mean values ranging from 88% (Cohort 4) to 94.4% (Cohort 6; **Figure 2C**). Nevertheless, medium-quality genomes can be used to reliably investigate abundance, prevalence, taxonomy, and genome size (Pacheco-Valenciana, Tausch, et al., 2025; Rodríguez-Gijón et al., 2025).

**Figure 2.**
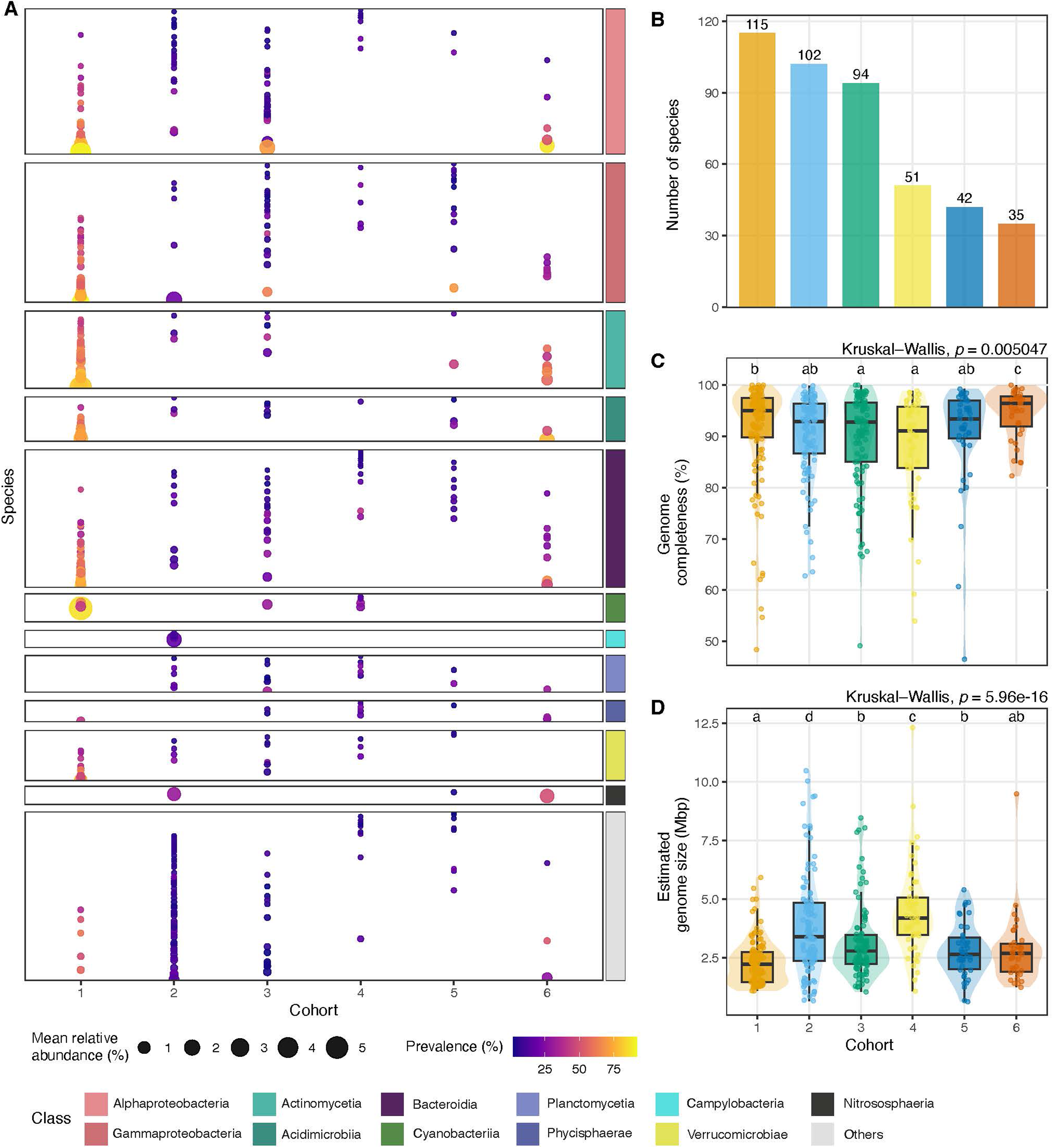
Taxonomic composition and genomic characteristics of Baltic Sea microbial cohorts. (A) Species composition of each cohort, grouped by taxonomy at the class level. Each row represents a species within a cohort, and a dot indicates species presence. Dot size indicates the mean relative abundance (%) across all samples, and the dot color gradient represents prevalence (%) across samples. Class identity is indicated by the different colors in the boxes. (B) Bars show the total number of species assigned to each cohort. Comparison of (C) genome completeness and (D) estimated genome size of species per cohort. Boxplots show the median and interquartile range (IQR), with whiskers extending to 1.5 x IQR. Statistical significance was assessed using a Kruskal–Wallis test, followed by Dunn’s post hoc test for pairwise comparisons. Groups sharing at least one letter (e.g., a and ab) are not significantly different, whereas groups with different letters (e.g., a vs. b) differ significantly (*p* < 0.05).

Cohorts differed in species richness (**Figure 2B)**, in their taxonomic composition and relative abundances and prevalences (**Figure 2A**). First, we observed that both Cohort 1 and Cohort 6 were overall composed of the most abundant and prevalent species (**Figure 2A** and **Supplementary Figure 1**). Species in these cohorts primarily belonged to class Alphaproteobacteria, Gammaproteobacteria, Actinomycetia, Acidimicrobiia, and Bacteroidia. Additionally, Cohort 1 contained abundant and prevalent Cyanobacteriia and Verrumicrobiae. In contrast, Cohort 2, which is associated with deeper, low-oxygen samples, was dominated by more diverse phylogenetic groups, including a species of Nitrososphaeria and four Campylobacteria, the latter being found only in this cohort. Cohort 3 showed a similar composition to Cohort 1 at the class level, but lacked strongly dominant species, whereas Cohorts 4 and 5 were characterized by generally lower abundances and fewer high-prevalence taxa across classes.

Across all species, mean relative abundance was positively associated with prevalence and, to a lesser extent, with the degree of connectedness in the co-occurrence network (**Supplementary Figure 2**). Residual distributions around these relationships varied among cohorts (**Supplementary Figure 3**). In the abundance–prevalence relationship (**Supplementary Figure 3A**), species in Cohort 1 tended to be more prevalent than expected based on their relative abundance. In contrast, for the prevalence–connectedness relationship (**Supplementary Figure 3B**), Cohort 2 included many species with positive residuals, indicating that several members were more connected than expected based on their prevalence. Estimated genome sizes showed pronounced differences among cohorts (**Figure 2D**). Cohort 1 had the smallest genomes on average (2.30 Mbp), followed by Cohorts 5 and 6, whereas Cohort 4 contained the largest average genomes (4.48 Mbp). Cohort 2 showed the second-largest average genome size (3.84 Mbp), and also the most variability. Notably, cohorts that are characterized by highly abundant and prevalent taxa (Cohorts 1 and 6) are also associated with the smallest estimated genome sizes and the highest coding density (**Supplementary Figure 4)**.

### A large-genome cohort with low oxygen preference contributes more functions for nitrogen and sulfur transformations

While high genome quality is required to robustly predict the functional potential of genomes, the high average completeness of genomes per cohort allows for comparative analyses of functional variation between cohorts. For this, we quantified the enrichment per cohort of genes involved in carbon, nitrogen, and sulfur transformations (**Figure 3**).

**Figure 3.**
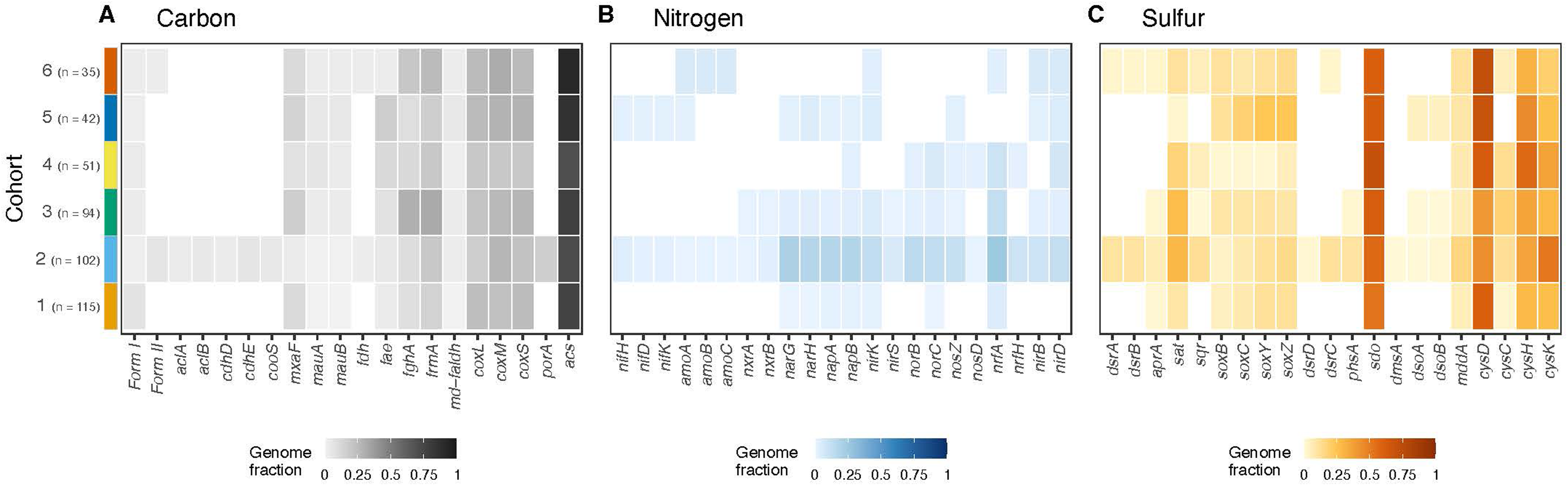
Enrichment of carbon, nitrogen, and sulfur genes across Baltic Sea microbial cohorts. Heatmaps show the proportion of genomes that have the gene present per cohort. Genes involved in the (A) carbon, (B) nitrogen, and (C) sulfur biogeochemical cycles across microbial cohorts, based on METABOLIC annotations. Rows correspond to cohorts, and columns to functional genes. Gene descriptions are provided in Supplementary Table 4. Color intensity indicates the fraction of genomes within each cohort that contain a given gene (0–1).

Carbon cycling genes, such as Rubisco Form I, were present across multiple cohorts (**Figure 3A)** in a relatively small fraction of genomes (2–7% across cohorts). The alternative carbon-fixation genes were more restricted and found only in a few genomes from Cohort 2. Genes for C1 metabolism, such as those for methanol, formaldehyde, and carbon monoxide oxidation, were detected in all cohorts, demonstrating their distribution among diverse microbial lineages in the Baltic Sea. The *porA* gene (pyruvate oxidation) was enriched exclusively in Cohort 2, present in 16.7% of genomes.

Nitrogen cycling genes exhibited the greatest differentiation among cohorts (**Figure 3B**). Cohort 2 contained the highest proportion of genomes with genes involved in various nitrogen processes, including nitrate reduction (*narG*, *narH*; 20.6% and 17.7% of genomes, respectively), periplasmic nitrate reduction (*napA*, *napB*; 17.7% and 18.6%), and nitric oxide reduction (*norB*, *norC*; 14.7% and 13.7%). Ammonia oxidation genes (*amoA*, *amoB*, and *amoC*) were mostly found in Cohorts 2 and 6, although these were restricted to one species in Cohort 2 and two species in Cohort 6. Nitrogen fixation genes were also rare and present only in a few genomes from Cohorts 2 and 5.

Sulfur metabolism genes were broadly distributed across cohorts (**Figure 3C**). Genes for thiosulfate oxidation (*soxBCYZ*) were found in all cohorts but were most prevalent in Cohort 5 where ∼20% of genomes encoded these genes, whereas Cohort 4 contained the smallest fraction (∼3%). Genes for dissimilatory sulfur metabolism (*dsrAB*) were limited to Cohorts 2 and 6. Genes involved in DMSO metabolism (*dsoAB*) appeared in one to two genomes in Cohorts 2, 3, and 5, and assimilatory sulfur metabolism (*cysCDHK*) genes were common, present in approximately 33% of genomes across all cohorts.

When considering species relative abundances (**Supplementary Figure 5**), these patterns became more pronounced, with alternative carbon fixation genes and nitrogen cycling functions mainly dominated by Cohort 2 taxa. Sulfur metabolism was mainly contributed by abundant members of Cohorts 2 and 6. These findings indicate that microbial cohorts partition biogeochemical functions, with specific cohorts disproportionately contributing to distinct carbon, nitrogen, and sulfur transformations.

### Three amino acids and three B-vitamins show the lowest pathway completeness across all cohorts

To explore biosynthetic potential across cohorts, we quantified the completeness of eighteen amino acid and nine B-vitamin biosynthesis pathways across genomes using custom pathway definitions based on KEGG modules (**Figure 4** and **Supplementary Figure 6**). Overall, we observed that, irrespective of genome quality, the capacity to biosynthesize both metabolite types was not equally retained. Most amino acid pathways were fully present in about 55% of genomes, across all pathways and cohorts. However, on average across cohorts, the pathways for methionine, phenylalanine, and tyrosine were rarely complete, occurring in only 11.9%, 5.1%, and 3.4% of genomes, respectively.

**Figure 4.**
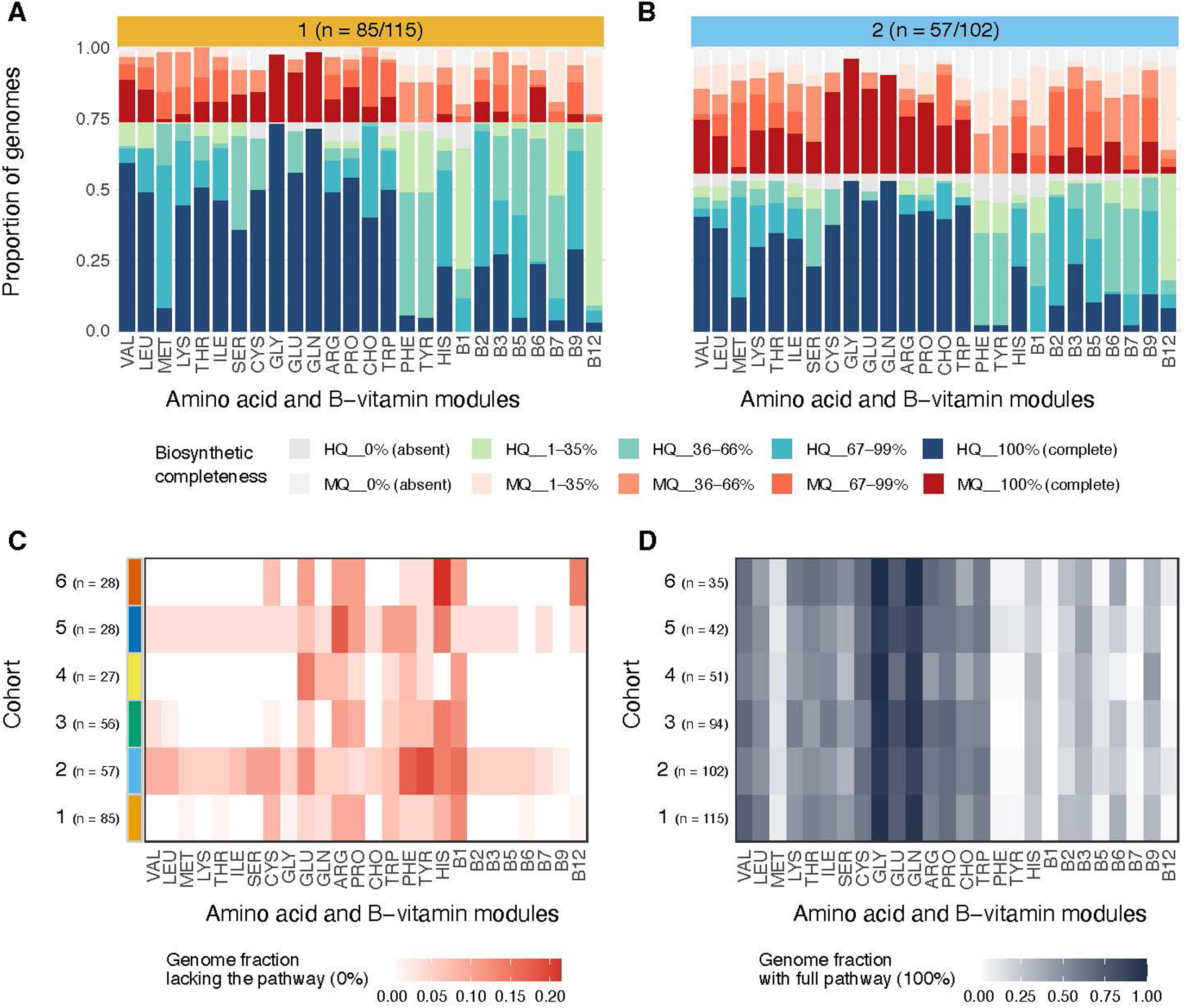
Amino acid and B-vitamin biosynthetic pathways across Baltic Sea microbial cohorts. Custom-defined module completeness for eighteen amino acid and nine B-vitamin biosynthesis pathways across members of (A) cohort 1 and (B) cohort 2. Species were grouped by genome quality into high-quality (HQ; >90% completeness and <5% contamination) and medium-quality (MQ; ≥50% completeness and <10% contamination) genomes. Stacked bar plots show the proportion of genomes across biosynthetic completeness categories for each pathway. Within each cohort, bars represent individual pathways, and the y-axis indicates the proportion of genomes assigned to each completeness group, further separated by genome quality (HQ at the bottom; MQ above). Biosynthetic completeness categories are defined as 0% (absent), 1–35%, 36–66%, 67–99%, and 100% (complete). For B_12_, completeness was defined as the best-route value, calculated as the higher of the aerobic and anaerobic custom module scores. Five color shades represent completeness levels for HQ genomes, and another five shades correspond to levels for MQ genomes. Numbers above each panel indicate the number of high-quality genomes relative to the total number of species within that cohort. (C) Heatmap showing the proportion of high-quality genomes per cohort that are fully lacking (0% completeness) a biosynthesis pathway. Rows represent cohorts (with number of species included in the analysis shown as n), and columns correspond to amino acid and B-vitamin biosynthesis pathways. Color intensity (white to red) indicates the fraction of genomes within each cohort with absent pathways. (D) Heatmap showing the proportion of genomes per cohort with a fully complete (100% completeness) biosynthetic pathway. Rows represent cohorts (with number of members in cohort shown as n), and columns correspond to amino acid and B-vitamin biosynthesis pathway. Color intensity (white to black) reflects the fraction of genomes with complete pathways.

The low completeness of methionine should be interpreted in the context of the specific module definition used here. In this analysis, methionine completeness was assessed using the *de novo* biosynthesis route represented by module M00017. The low methionine pathway completeness was mainly driven by the frequent absence of *metB* across cohorts. In contrast, *metE* and *metH*, which catalyze the final conversion of L-homocysteine to methionine, were common across cohorts.

Similarly, the biosynthetic potential for phenylalanine and tyrosine should be interpreted with caution. In our custom modules, these pathways start at chorismate, the main precursor for aromatic amino acid biosynthesis. The chorismate biosynthesis pathway was analyzed separately and was 100% complete in 48% of genomes across cohorts on average, while an additional 40% of the genomes included 67–99% of the pathway. Thus, the low full completeness of phenylalanine and tyrosine mainly reflects that few genomes encoded all downstream steps from chorismate to the final amino acid. Because these branches consist of only three steps, many genomes still retain partial pathway capacity, so the apparent absence of pathways could reflect either potential metabolic dependence or limitations in annotations, as alternative or poorly characterized enzymes may not be captured by standard KEGG modules.

On the other hand, B-vitamin biosynthesis pathways were more often partially present. In particular, for vitamins B_1_, B_7_, and B_12_, very few genomes encoded either the full pathway (∼3.3%) or lacked the pathway entirely (∼8.7%). Most genomes contained partial pathways. For vitamin B_1,_ most of the genomes (43.5%) had a pathway completeness between 1 and 35%. For vitamin B_12_, 76.5% of the genomes contained less than 35% of the pathway, whereas vitamin B_7_ showed the highest proportion of genomes (46.2%) with medium pathway completeness (36–66%).

When examining the proportions of genomes with either full presence or absence of biosynthesis pathways across all cohorts, clear differences between cohorts emerged (**Figure 4C** and **D**). There are some cohorts for which very few genomes fully lack the biosynthesis pathway for a metabolite (Cohorts 1, 3, 4, and 6; **Figure 4C**). In contrast, Cohorts 2 and 5 showed the largest fraction of genomes lacking several biosynthetic pathways. As for the genomes with complete pathways (**Figure 4D**), the pattern remains: amino acids have the largest proportion of genomes with full pathways, and vitamins have the smallest. These patterns were consistent when comparing analyses restricted to high-quality genomes with those including all genomes (**Supplementary Figure 7**), indicating that the observed distributions of absent (0%) and complete (100%) pathways were largely preserved.

### Most cohort members have B_12_ biosynthesis genes but the full pathway is contained in a few low abundance Alphaproteobacteria

Because B-vitamins frequently occur as incomplete pathways across genomes, we focused on B_12_ biosynthesis as a representative case. Vitamin B_12_ has a large pathway that would allow us to observe nuances even in medium-quality genomes. The biosynthesis of vitamin B_12_ involves two possible corrin ring biosynthesis pathways: an anaerobic route (M00924) and an aerobic route (M00925), each with 11 module steps that lead to the production of Cob(II)yrinate a,c diamide. In this framework, some enzymatic transformations are merged into single module steps, which is why the number of module steps does not exactly match all individual biochemical reactions. The cobalamin precursor is finally modified through the nucleotide loop assembly pathway (M00122), which includes corrinoid remodeling and lower-ligand activation steps, ultimately resulting in B_12_ synthesis. However, our analysis does not by itself indicate *de novo* lower-ligand biosynthesis or identify the final lower ligand.

In our dataset, very few genomes encoded the complete B_12_ biosynthesis pathway, but only a small fraction lacked it entirely (**Figure 4C** and **D**), resulting in a large proportion of genomes containing a partial pathway. To examine how these steps are distributed across microbial cohorts, we analyzed the proportion of genomes encoding each gene in these modules (**Figure 5**). We observed that genes belonging to the nucleotide loop assembly pathway (M00122) were more common, especially its first step (*cobA*), which catalyzes the adenosylation step and was highly prevalent in most cohorts (nearly universal, 80 to 88% of genomes) (**Figure 5C**).

**Figure 5.**
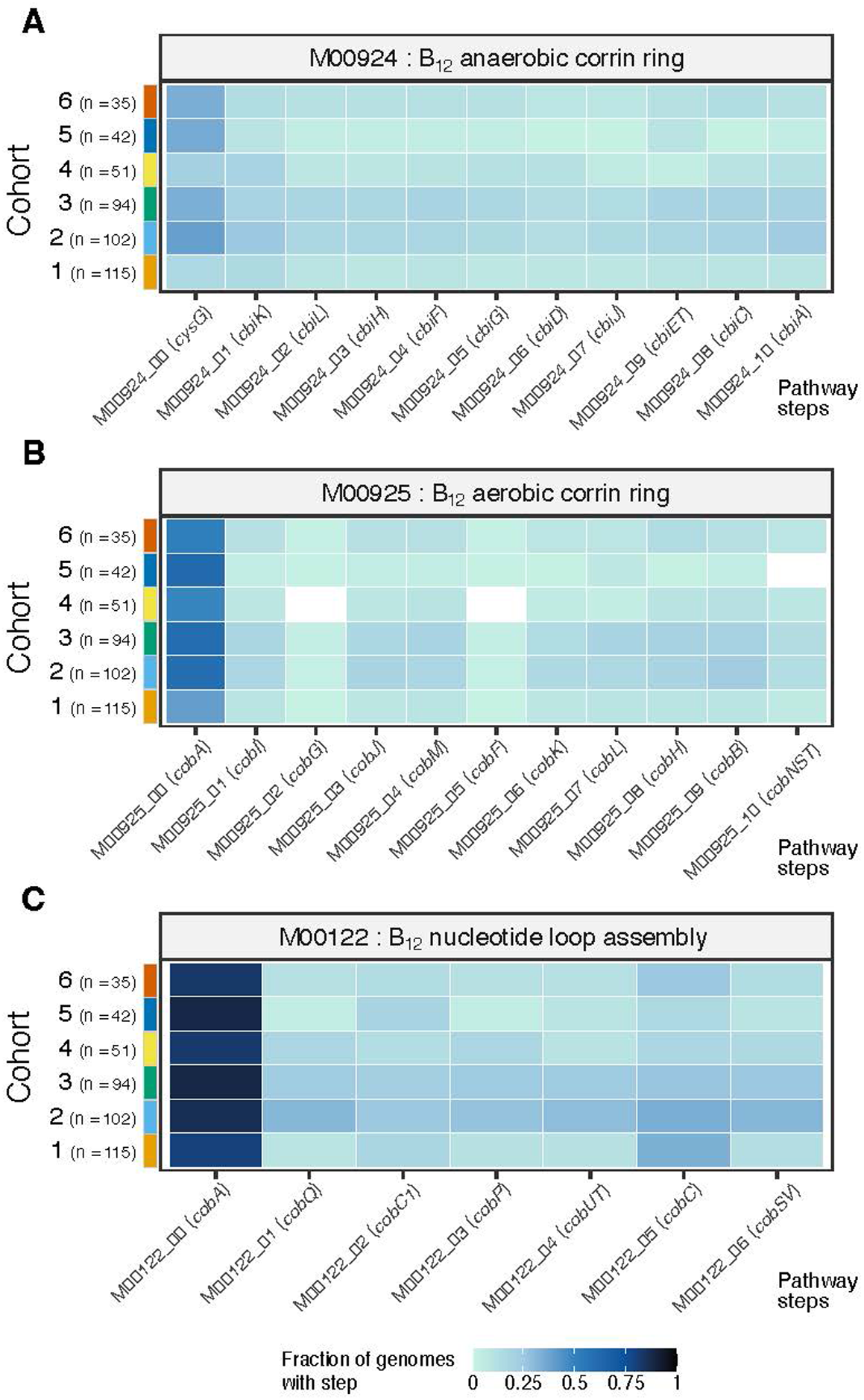
Distribution of vitamin B_12_ biosynthetic modules across Baltic Sea microbial cohorts. Heatmaps showing the gene-level distribution of vitamin B_12_ biosynthesis modules across microbial cohorts. (A) M00924: anaerobic corrin ring biosynthesis pathway. (B) M00925: aerobic corrin ring biosynthesis pathway. (C) M00122: nucleotide loop assembly. Rows represent cohorts (cohort size indicated as n), and columns correspond to individual genes within each module. Color intensity reflects the fraction of genomes per cohort encoding each gene (0–1).

To investigate the distribution of B_12_ biosynthetic steps at higher resolution, we analyzed gene presence across all species within Cohort 1 (the largest cohort with the most high-quality genomes, **Figure 6A**). The corrin ring biosynthesis pathways (M00924 or M00925) were distributed across a few genomes within Alphaproteobacteria and Cyanobacteriia. Alphaproteobacterial genomes carrying these genes also encoded the full nucleotide loop assembly pathway, whereas Cyanobacteriia retained only a subset of nucleotide loop assembly genes, mostly those associated with corrinoid remodeling, and lacked the lower-ligand activation genes. Among all other genomes, genes from the nucleotide loop assembly pathway were more consistently retained. Comparisons of module completeness confirmed that the nucleotide loop assembly pathway was significantly more complete across genomes than either corrin ring biosynthesis pathway (**Figure 6B**; Kruskal–Wallis, *p* = 2.2e-16). This view revealed taxonomic patterns in the distribution of individual pathway steps within the nucleotide loop assembly pathway. For instance, some genes were more frequently detected within specific classes, such as *cobC1* in Gammaproteobacteria and *cobC* in Actinomycetia, indicating taxonomic differences in the distribution of downstream nucleotide loop assembly reactions.

**Figure 6.**
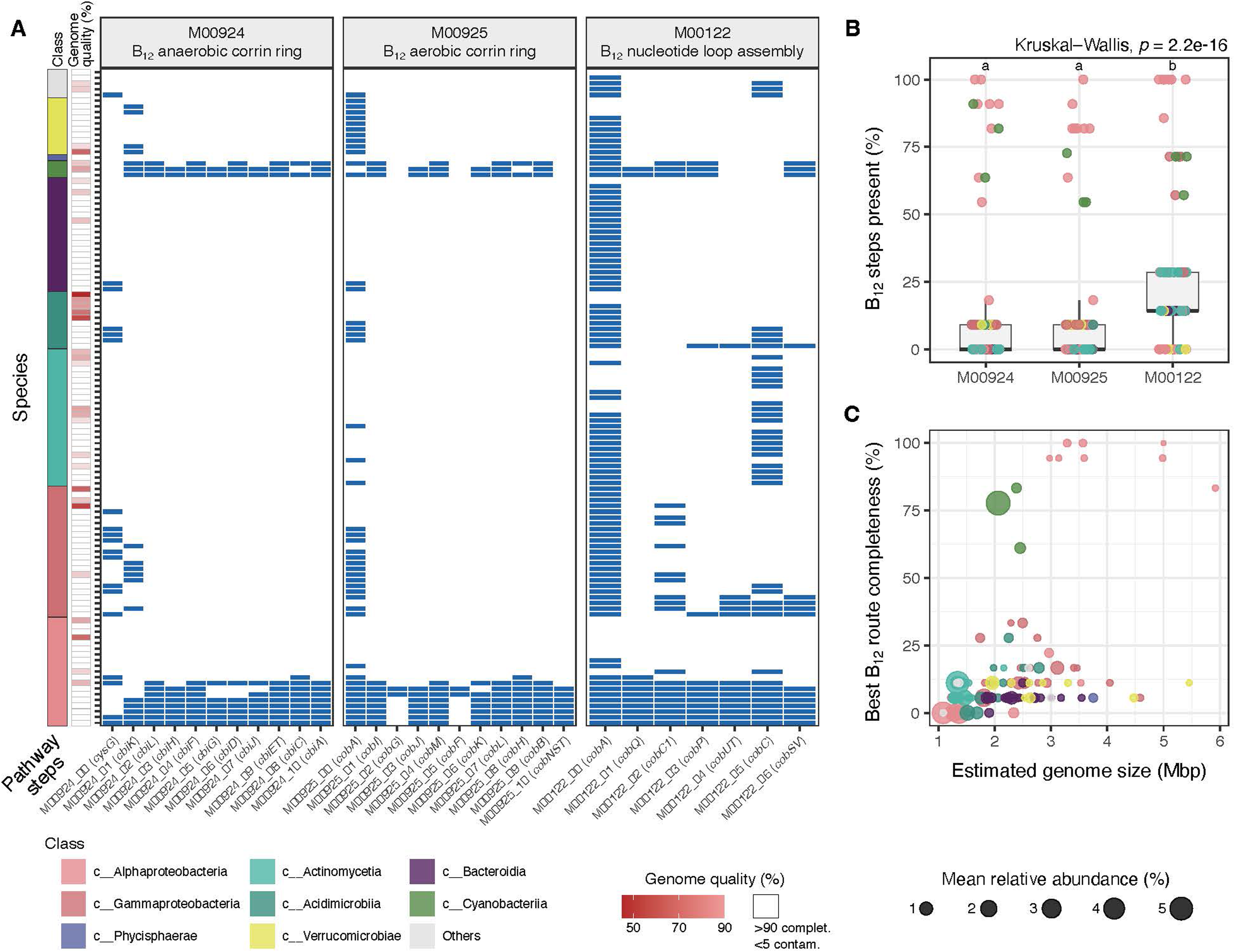
Gene-level distribution of vitamin B_12_ biosynthetic pathways in Cohort 1 and links to ecological traits. (A) Tile plot showing the presence or absence of genes associated with three vitamin B_12_ biosynthetic modules across all species in Cohort 1: M00924 (anaerobic corrin ring pathway), M00925 (aerobic corrin ring pathway), and M00122 (nucleotide loop assembly). Rows represent individual species (n = 115), ordered by taxonomic class, and columns correspond to individual pathway steps (genes) within each module. Blue tiles indicate gene presence; white tiles indicate absence. All genomes are shown regardless of quality. The side annotation denotes genome quality: white indicates high-quality genomes (>90% completeness and <5% contamination), whereas red shading indicates genomes below these thresholds, with darker red corresponding to lower completeness. **(B)** Boxplots showing within-module completeness of B_12_ biosynthesis across species. For each species and module, the proportion of encoded steps was calculated relative to the total number of steps in that module. Boxplots show the median and interquartile range (IQR); whiskers extend to 1.5 × IQR; points represent individuals. Differences among modules were evaluated using a Kruskal–Wallis test followed by Dunn’s post hoc test for pairwise comparisons. Modules sharing a letter are not significantly different, whereas modules with distinct letters differ significantly (*p* < 0.05). **(C)** Correlations between best-route B_12_ biosynthetic completeness and genome size across species in Cohort 1. For each species, completeness was calculated separately for the anaerobic route (M00924 + M00122) and the aerobic route (M00925 + M00122), and the higher value was retained as the best-route completeness. Estimated genome size (Mbp) is shown on the x-axis and best-route completeness on the y-axis. Point size reflects the mean relative abundance across samples.

We also examined how the completeness of the B_12_ biosynthesis pathway relates to genome size and relative abundance (**Figure 6C**). On the one hand, genomes with pathway completeness above 85% were mostly larger than 3 Mbp, had lower abundance (<0.12%), and all belonged to Alphaproteobacteria. On the other hand, the most abundant species (>3%) were associated with smaller genomes (∼1–1.4 Mbp) encoding only up to 17% of B_12_ biosynthetic steps, particularly among Alphaproteobacteria, Gammaproteobacteria, Actinomycetia, and Acidimicrobiia. Cyanobacteriia stood out as an exception to this pattern because they had comparatively small genomes (between 2.06 and 2.45 Mbp), high relative abundance (up to 5%), and completeness of the B_12_ biosynthesis pathway between 61 and 83%.

### Biosynthetic pathways with low completeness are taxonomically partitioned within cohorts

The B_12_ case study showed that biosynthetic potential can be unevenly distributed across species within cohorts, with only a few taxa encoding the full pathway, while most genomes retained only partial pathways. To assess whether this pattern extends beyond B_12_, we next examined the six metabolites with the lowest pathway completeness across cohorts (methionine, phenylalanine, tyrosine, and vitamins B_1_, B_7_, and B_12_) and how their biosynthetic potential is distributed across taxonomic groups (**Figure 7**).

**Figure 7.**
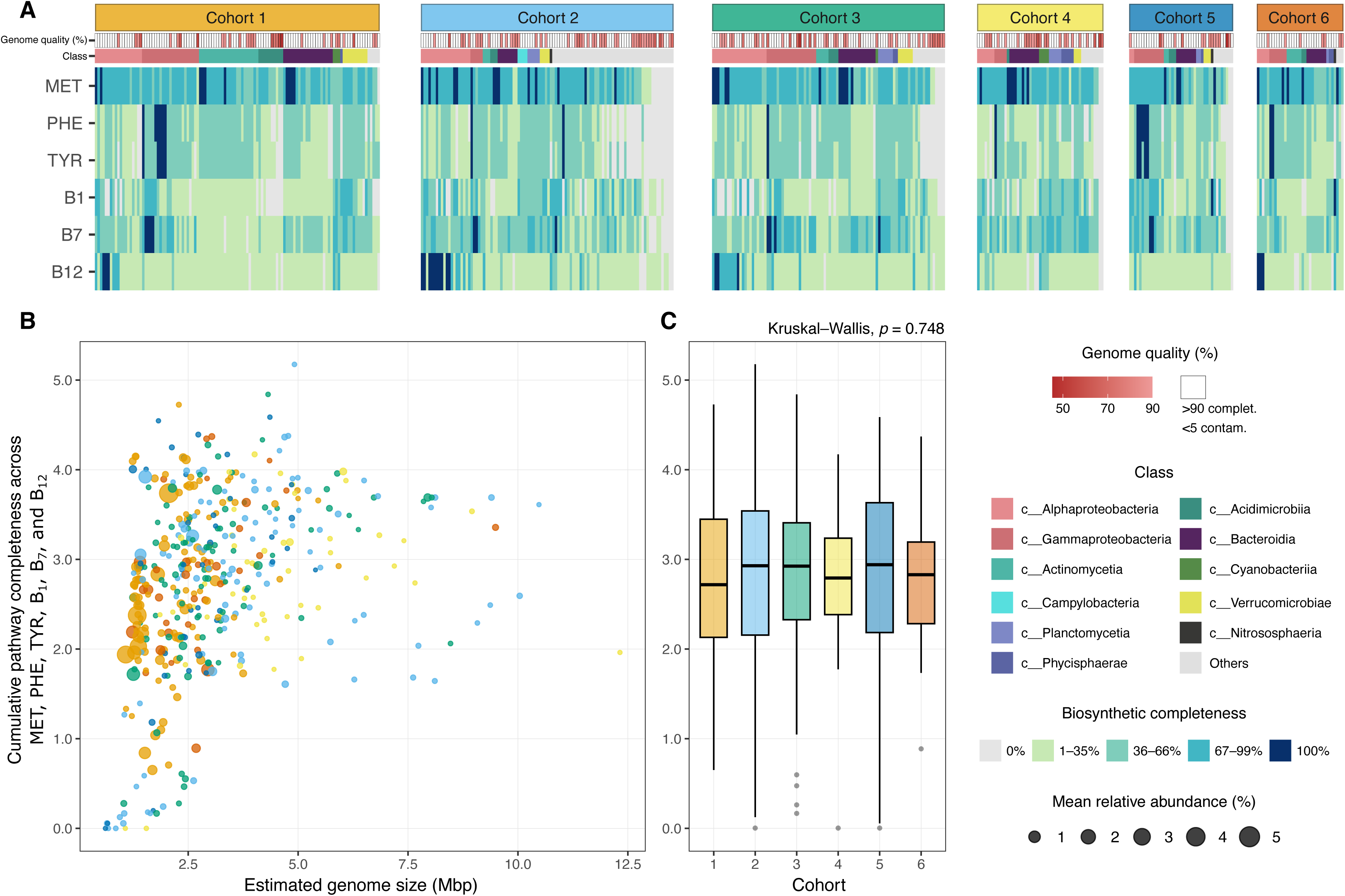
Patterns of biosynthetic pathway completeness for selected metabolites across species within Baltic Sea microbial cohorts. (A) Heatmap showing the biosynthetic pathway completeness for the six metabolites with the lowest proportion of genomes encoding fully complete (100%) pathways across species from Cohorts 1–6. Columns represent individual species within each cohort, ordered by taxonomic class and cumulative completeness across the six selected metabolites. Rows correspond to the selected metabolites. Tile colors indicate biosynthetic completeness categories: 0% (absent), 1–35%, 36–66%, 67–99%, and 100% (complete). For B_12_, completeness was defined as the best-route value, calculated as the higher of the aerobic and anaerobic custom module scores. Top annotation bars indicate genome quality and taxonomic class. High-quality genomes are defined as >90% completeness and <5% contamination. (B) Scatter plot showing the relationship between estimated genome size (Mbp) and cumulative pathway completeness across the same six selected metabolites, including three amino acids (methionine, phenylalanine, and tyrosine) and three B-vitamins (B_1_, B_7_, and B_12_), for species belonging to Cohorts 1–6. Cumulative pathway completeness was calculated as the sum of module completeness values across the six metabolites (range: 0–6). Each point represents one species. Point size indicates mean relative abundance (%) across samples, and point color indicates cohort assignment. (C) Boxplots show the distribution of cumulative pathway completeness across species within each cohort, using the same cumulative completeness metric as in panel B. Boxplots display the median and interquartile range (IQR), and whiskers extend to 1.5 × IQR. Differences among cohorts were tested with a Kruskal–Wallis test.

Across all cohorts, these six metabolites showed clear taxon-specific differences rather than a uniform distribution across the community. Some pathways were strongly associated with particular classes. Across cohorts, B_12_ biosynthesis remained 100% complete, almost entirely within Alphaproteobacteria, while several other groups showed only low pathway completeness (1–35%), including Gammaproteobacteria, Actinomycetia, Bacteroidia, Planctomycetia, and Verrucomicrobiae. In contrast, complete biosynthetic pathways for the aromatic amino acids phenylalanine and tyrosine were mainly associated with Gammaproteobacteria, while several other taxa showed only partial representation of these pathways. Patterns of pathway absence and presence also revealed differentiation within taxonomic groups. Gammaproteobacteria, for example, largely lacked the capacity for B_12_ biosynthesis, even though they dominated the signal for phenylalanine and tyrosine, and contributed to B_7_ biosynthesis through different species within the class.

Across species, cumulative completeness across these six metabolites remained limited, with larger genomes often showing higher cumulative completeness values, whereas several smaller but relatively abundant genomes showed intermediate values (**Figure 7B**). However, the overall distribution of cumulative completeness was similar across cohorts and did not differ significantly among them (**Figure 7C**; Kruskal–Wallis, *p* = 0.748).

Methionine followed a different pattern. Although complete *de novo* methionine biosynthesis was still uncommon, near-complete pathways were widespread and distributed across almost all classes in our dataset. Approximately 71% of genomes encoded near-complete methionine pathways, while only ∼11.9% encoded the complete *de novo* methionine biosynthesis pathway. To further resolve the relationship between methionine synthesis and B_12_ biosynthetic potential, we examined the distribution of the methionine synthase genes *metE* and *metH* across genomes grouped by the B_12_ biosynthesis completeness. This comparison is relevant because *metH* mediates B_12_-dependent methionine synthesis, whereas *metE* encodes a B_12_-independent alternative (**Supplementary Figure 8**). We observed that 67–70% of genomes with low B_12_ biosynthesis pathway completeness (1–35%) encoded *metH* without the B_12_-independent alternative *metE*, whereas genomes with increasing B_12_ completeness showed a progressive shift toward the combined presence of both *metH* and *metE*. In genomes with complete B_12_ biosynthesis pathways, most encoded both enzymes (61–67%), while the fraction encoding *metH* alone decreased substantially. This pattern was consistent across the full Baltic Sea MAG dataset (n = 701) and the high-quality subset (n = 450), indicating that it is not driven by genome incompleteness.

Taxonomic associations and metabolite biosynthesis were broadly conserved across cohorts, although some variability remained within particular class–metabolite combinations. Overall, the main signal was not that each cohort had a completely different biosynthetic profile, but that different metabolites were repeatedly associated with different taxonomic groups. Methionine stood out as an exception to this broader pattern because, despite its low full *de novo* pathway completeness, near-complete pathways were common, and the distribution of *metE* and *metH* revealed a widespread potential for B_12_-dependent methionine synthesis. These results indicate that biosynthetic potential is strongly partitioned across taxa within Baltic Sea microbial cohorts, reinforcing metabolic dependencies.

## DISCUSSION

Microbial communities are more than just random collections of individual species. They are ecologically organized groups whose traits result from the combined characteristics of the populations living together (Faust & Raes, 2012; Fuhrman et al., 2015; Widder et al., 2016). They have also been proposed to function as metabolically interconnected systems shaped, at least in part, by metabolic dependencies among their members (Zelezniak et al., 2015). In our study, we further investigated the system interconnectedness by analyzing the potential catabolism and anabolism of co-occurring groups of microorganisms in the Baltic Sea. These microbial cohorts are not closed systems and may also interact with other surrounding microbes or abiotic factors. Nevertheless, our cohort-based analysis shows that Baltic Sea microbial cohorts differ consistently in their environmental distributions, genomic traits, and catabolic and biosynthetic potential.

However, genome-resolved metagenomics can only describe the fraction of the microbial community represented in the underlying genome catalog, so our cohort analysis should be interpreted within that constraint. In our dataset, the Baltic Sea species catalog captured a substantial but incomplete portion of the Baltic Sea microbiome, and the mapped fraction varied across the salinity gradient from 5 to 64%. This range is similar to that of the best representing marine genome collection (Paoli et al., 2022). Even so, the represented fraction was ecologically informative, and these cohorts recovered clear environmental structure across the Baltic Sea. This suggests that the inferred cohorts are meaningful despite potentially missing some members.

Inferring co-occurrence networks from abundance data is data-intensive, and similar analyses based on amplicon data indicate that while cohort structure is robust to undersampling, the total number of cohort members increases with sample size (Milke et al., 2026). Nevertheless, even though 16S-based amplicon studies typically report higher diversities per sample, previously characterized amplicon-based aquatic cohorts contained between 10 and 300 members (Milke et al., 2026), similar to our findings. While the total diversity within cohorts likely increases with MAG catalogs of higher diversity representation, the consistent patterns observed indicate that our framework captures robust and ecologically meaningful organization of the Baltic Sea microbiome.

One clear pattern was the relationship between genome size, abundance, prevalence, and connectedness in the co-occurrence network. Members of Cohort 1 contained the smallest genomes and included many of the most abundant and prevalent taxa, yet their connectivity in the co-occurrence network was, on average, lower than that of Cohort 2. Taxa in Cohort 2 appeared more connected than their prevalence alone would suggest. This differs from freshwater observations, where abundant and prevalent cohorts represented by streamlined taxa were also strongly connected (Rodríguez-Gijón et al., 2025). Our results therefore suggest that, in the Baltic Sea, ecological success, prevalence, and connectedness do not necessarily align across cohorts.

Functional differentiation among cohorts was also apparent in the distribution of biogeochemically relevant genes. The clearest example was Cohort 2, which was associated with deeper and lower- oxygen conditions and was enriched in genes involved in carbon fixation, nitrogen transformations, and sulfur-related functions. This pattern agrees with earlier work showing that microbial community composition and functional traits shift sharply across depth and redox gradients in the Baltic Sea, with deeper waters showing stronger representation of nitrogen and sulfur cycling processes (Thureborn et al., 2013). More broadly, Baltic Sea microbial communities and their functions are known to be strongly structured by salinity and physicochemical heterogeneity (Dupont et al., 2014; Herlemann et al., 2011), and our cohort-based framework recovers similar ecological patterns at the level of recurring groups of co-occurring species. The important point here is not that every member of a cohort contributes equally to these functions, but that some functions are disproportionately represented in specific cohorts. In that sense, cohorts are environmentally filtered subcommunities with selected metabolic repertoires.

Beyond catabolic traits linked to environmental gradients, a clear anabolic pattern we found was the difference between amino acid and B-vitamin biosynthesis. Across cohorts, amino acid biosynthesis pathways were more often complete, while B-vitamin biosynthesis pathways were rather incomplete or patchy. This aligns with genomic studies showing loss of metabolic functions leading to increased metabolic dependencies (D’Souza et al., 2014; Giordano et al., 2024; Giovannoni et al., 2014; Kost et al., 2023; Rodríguez-Gijón et al., 2025). The Black Queen Hypothesis proposes that biosynthetic functions can be lost in community members as long as the producers supply the necessary amounts of such metabolites to support the community (Morris et al., 2012). Amino acids are indispensable building blocks for protein synthesis, which likely encourages the retention of complete biosynthetic pathways. B-vitamins function as enzyme cofactors and are therefore required in much smaller amounts, or in some prokaryotes, not even required at all (Gregor et al., 2025; Groon et al., 2025; Sañudo-Wilhelmy et al., 2014). This lower cellular demand may make maintaining the full biosynthetic pathway less favorable (Helliwell et al., 2013; Shelton et al., 2019).

Vitamin B_12_ provides a particularly informative case study for this broader pattern. In our dataset, very few genomes encoded the complete B_12_ biosynthesis pathway, yet many retained at least one gene associated with it. Genes from the nucleotide loop assembly pathway were more common than those from corrin ring biosynthesis, showing an uneven distribution of B_12_ biosynthesis across taxa. In Cohort 1, only three Alphaproteobacterial genomes encoded a complete *de novo* B_12_ biosynthesis pathway, while five additional Alphaproteobacterial genomes encoded an almost complete pathway (∼90%) with the full nucleotide loop assembly pathway but missing some corrin ring biosynthesis genes. In addition, only three Cyanobacteriia encoded a near-complete pathway. Thus, genomes with near-complete to complete corrinoid biosynthesis represented only a small fraction of Cohort 1, whereas many more retained genes from the nucleotide loop assembly pathway. This restricted distribution matches known ways in which microorganisms depend on each other by salvaging corrinoid ring intermediates or remodeling unsuitable cobamides (B_12_-analogues with a different or unsuitable lower ligand attached), rather than synthesizing the entire pathway *de novo*, relying on external supply of corrinoids, cobamides, lower ligands, or related intermediates (Alvarez-Aponte et al., 2025; Shelton et al., 2019; Villa & Escalante-Semerena, 2024; Wienhausen et al., 2024).

Cyanobacteriia are particularly interesting in this context because many are known to produce pseudocobalamin (with adenine rather than DMB as the lower ligand), a cobamide that is less bioavailable to most organisms (Heal et al., 2017; Helliwell et al., 2016). However, our analysis does not resolve *de novo* lower-ligand biosynthesis or the identity of the lower ligand, and therefore any inference about whether the Cyanobacteriia in our dataset produce pseudocobalamin or another cobamide variant should be made with caution. Instead, these genomes retained only a subset of nucleotide loop assembly genes, primarily those associated with corrinoid remodeling and downstream assembly. Nonetheless, this raises the possibility that Cyanobacteriia contribute pseudocobalamin-like corrinoids that may then be remodeled by other cohort members. Because B_12_ and its respective intermediates can influence microbial growth, interactions, and community assembly in aquatic systems (Bertrand et al., 2007; Browning et al., 2017; Croft et al., 2005; Gregor et al., 2025; Wienhausen et al., 2022), the uneven distribution of biosynthetic steps observed in this study is likely to have ecological consequences. B_12_ biosynthesis may nevertheless be maintained at the community level, as different taxonomic groups can contribute to corrinoid production under varying environmental conditions, providing a form of functional redundancy within microbial communities (Beauvais et al., 2023).

A supportive line of evidence comes from the distribution of methionine synthesis strategies across B_12_ biosynthesis pathway completeness categories. Genomes with low B_12_ biosynthesis pathway completeness were dominated by *metH*, lacking the B_12_-independent alternative *metE*, whereas genomes with increasing B_12_ biosynthesis completeness showed a progressive shift toward the combined presence of *metH* and *metE*. This pattern suggests that taxa with limited B_12_ biosynthetic potential may more often rely on B_12_-dependent methionine synthesis, whereas taxa with higher biosynthetic capacity tend to retain both methionine synthase genes. This aligns with previous observations that loss of the B_12_-independent methionine synthase *metE* is associated with the evolution of B_12_ dependence (Helliwell et al., 2015), and suggests that a similar pattern may extend across Baltic Sea genomes. More broadly, this interpretation is consistent with large-scale genomic surveys showing that cobamide producers represent only a subset of microbial communities, whereas potential cobamide users are much more widespread, with *metH* among the most commonly recovered cobamide-dependent functions in both marine and soil genomes (Shelton et al., 2019; Wang et al., 2024). In contrast, retention of both methionine synthases may allow greater flexibility in response to changing B_12_ availability (Groon et al., 2025; Rao et al., 2024). The retention of *metH* in genomes with low B_12_ biosynthetic potential is also consistent with the idea that some taxa may forgo the cost of complete corrinoid biosynthesis while retaining the capacity to use externally supplied B_12_ (Morris et al., 2012). At the same time, B_12_ should not necessarily be assumed to be the main metabolite connecting the members of each cohort, since these taxa may also access corrinoids produced by members of co-occurring cohorts, and we found biosynthetic pathways for other metabolites to be absent across all species in the Baltic Sea.

However, the taxonomically structured distribution observed for vitamin B_12_ reflects a broader organization of biosynthetic potential. Likewise, other metabolites with low pathway completeness also showed structured distributions across taxa. Rather than being randomly scattered across the community, biosynthetic potential for these metabolites was repeatedly associated with particular taxonomic groups, while other groups consistently lacked the same pathways (**Figure 7**). Because these taxa co-occur within the same cohorts, this pattern suggests that co-existing members contribute different subsets of anabolic functions. Importantly, this does not imply direct metabolic exchange between taxa. Instead, metabolites and their intermediates may also become available through dissolved environmental pools generated by microbial mortality and viral lysis, which release intracellular compounds into the surrounding environment and can thereby support organisms lacking complete biosynthetic pathways (Breitbart et al., 2018; Moran et al., 2022; Pande & Kost, 2017; Sultana et al., 2025).

Overall, our results indicate that environmental filtering and metabolic potential are jointly reflected in the organization of Baltic Sea microbial cohorts. The cohort framework captures groups of co-occurring species whose genomic repertoires align with salinity, depth, and oxygen-related conditions. In addition to these environmental associations, we observed a consistent contrast across cohorts between broadly retained amino acid biosynthesis pathways and B-vitamin biosynthesis pathways, the latter being restricted to relatively few members, as well as a non-random distribution of biosynthetic capabilities across taxa within cohorts. These patterns indicate that microbial cohorts capture ecologically structured assemblages in which anabolic and catabolic functions are unevenly distributed among co-existing members, and suggest that B-vitamins, more than amino acids, may represent a stronger axis of metabolic interdependence within cohorts. At the same time, these conclusions remain bounded by the current representation of the Baltic Sea microbiome in the genome catalog. Additional sampling, improved genome recovery, and integration with transcriptomic, metabolomic, and experimental approaches will be needed to determine whether the metabolic interdependencies suggested by these genomic patterns are actively realized in situ. Nonetheless, the present results show that microbial cohorts provide a useful ecological framework for linking environmental gradients, genomic traits, and the partitioning of anabolic and catabolic functions across natural microbial communities.

## Data Availability

All metagenomes and the genome catalog used in this study are publicly available from previously published studies (Pacheco-Valenciana, Garcia, et al., 2025; Pacheco-Valenciana, Tausch, et al., 2025). Accession numbers are PRJNA1134408, PRJEB22997, PRJEB34883, and SRP077551 (**Supplementary Table 2**).

## Code Availability

The code used in this study is available on GitHub at https://github.com/armandopv/baltic-sea-microbial-cohorts-metabolism. The repository includes all scripts and input tables required to reproduce the analyses and figures.

## ACKNOWLEDGMENTS

This work was funded by SciLifeLab and by the Swedish Research Council VR (grant 2022-03077). The authors acknowledge support from SNIC/Uppsala Multidisciplinary Center for Advanced Computational Science for access to the UPPMAX computational infrastructure, as well as the National Academic Infrastructure for Supercomputing in Sweden (NAISS). Computational work and data handling were enabled by resources in the projects SNIC 2022/5-392, 2023/5-126, NAISS 2023-5-379, NAISS 2024/6-411, NAISS 2024/5-156, NAISS 2025/22-777, and NAISS 2025/6-182 provided by the Swedish National Infrastructure for Computing (SNIC) at UPPMAX, and the National Academic Infrastructure for Supercomputing in Sweden (NAISS) partially funded by the Swedish Research Council through grant agreement 2018-05973 and 2022-06725, respectively. Support for this work was also provided by DFG grant (project number: 550295176). The authors acknowledge the use of ChatGPT (OpenAI) for language editing and clarity improvement.

## AUTHOR CONTRIBUTIONS

S.L.G. conceptualized and supervised the research. A.P.-V., F.M., and G.W refined the research idea. A.P.-V. and F.M. performed the bioinformatics analysis. A.P.-V. selected and performed statistical tests. A.P.-V. and F.M. performed the data interpretation and visualization. A.P.-V. led the writing of the manuscript with input from S.L.G. All co-authors contributed to the literature search, participated in editing and reviewing the manuscript, and approved the final version.

## Notes

https://github.com/armandopv/baltic-sea-microbial-cohorts-metabolism

### Competing Interest Statement

The authors have declared no competing interest.

