## Supplementary Figures 1 to 8 for "Baltic Sea microbial cohorts exhibit catabolic specialization and anabolic interdependencies across environmental gradients"

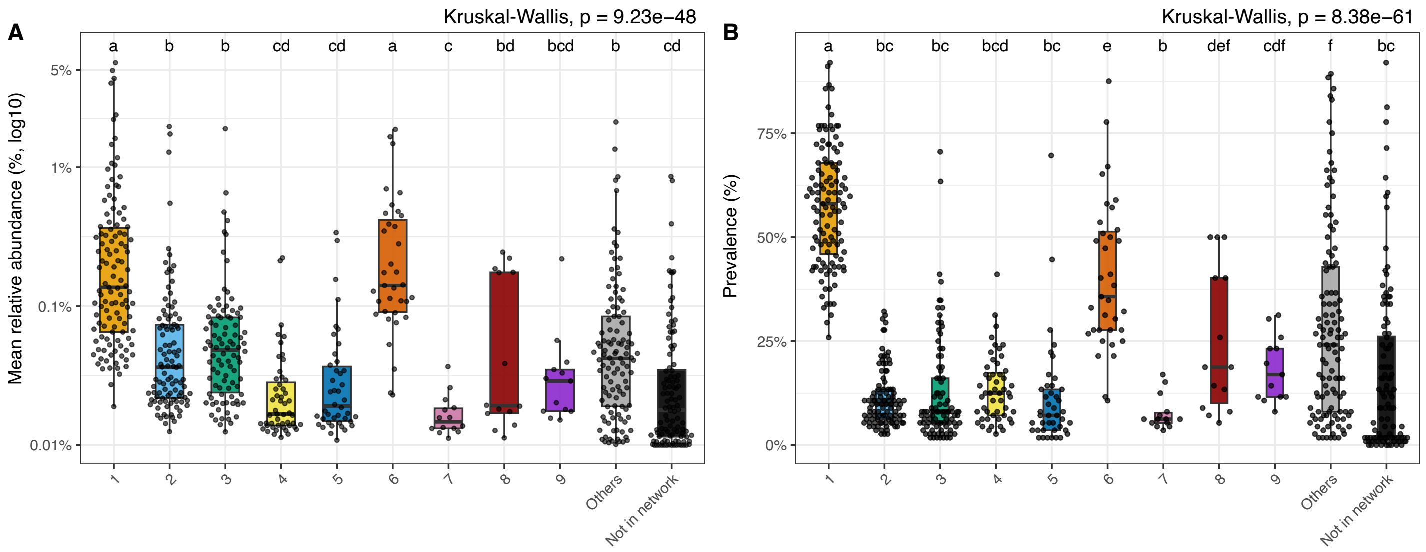


**Supplementary Figure 1. Mean relative abundance and prevalence of cohort members.** Boxplots show the distribution of (a) mean relative abundance (%, log10) and (b) prevalence (%) of species clusters across cohorts, as well as species classified as "Others" and "Not in network". Overall differences among cohorts were tested using a Kruskal–Wallis test, followed by Dunn's post hoc test for pairwise comparisons. Letters above boxplots denote significance groups ($P < 0.05$); groups sharing at least one letter are not significantly different. Boxplots show the median and interquartile range (IQR), with whiskers extending to 1.5× IQR; points represent individual species.

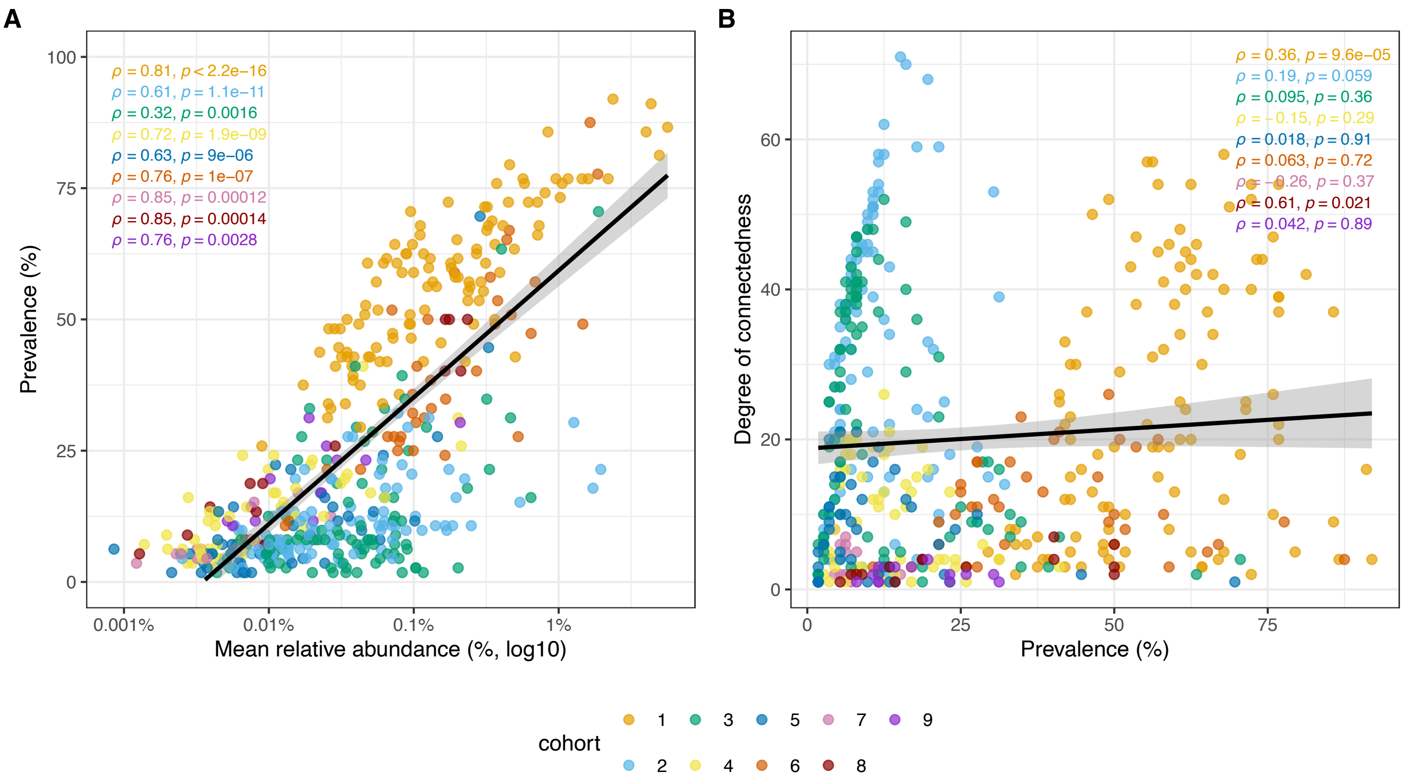


**Supplementary Figure 2. Relationship between relative abundance, prevalence, and degree of connectedness of cohort members.** Scatter plots showing the relationship between (A) mean relative abundance (%, log10) and prevalence (%) and (B) prevalence and degree of connectedness (%) for all species assigned to cohorts (1–9). Each point represents a species and is colored by cohort. Spearman correlations between mean relative abundance and prevalence or connectedness were computed for cohort members within each cohort (ρ and P values shown). A linear regression line fitted across all species clusters is shown for visual guidance only.

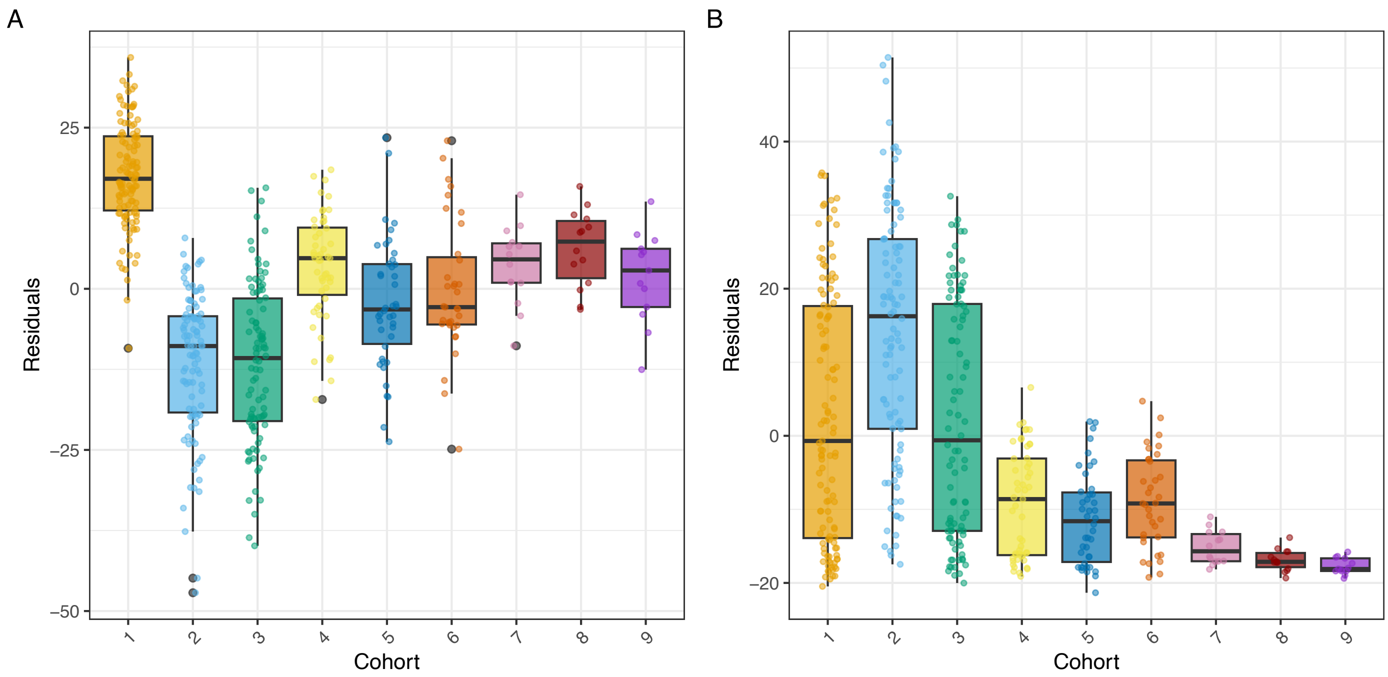


**Supplementary Figure 3. Cohort-specific residuals from abundance and prevalence relationships.** Boxplots showing the distribution of residuals from linear models relating (A) mean relative abundance to prevalence (%) and (B) prevalence to degree of connectedness across all species per cohort. Residuals indicate deviations of individual species from the global regression shown in Supplementary Figure 2. Points represent individual species grouped by cohort (1–9).

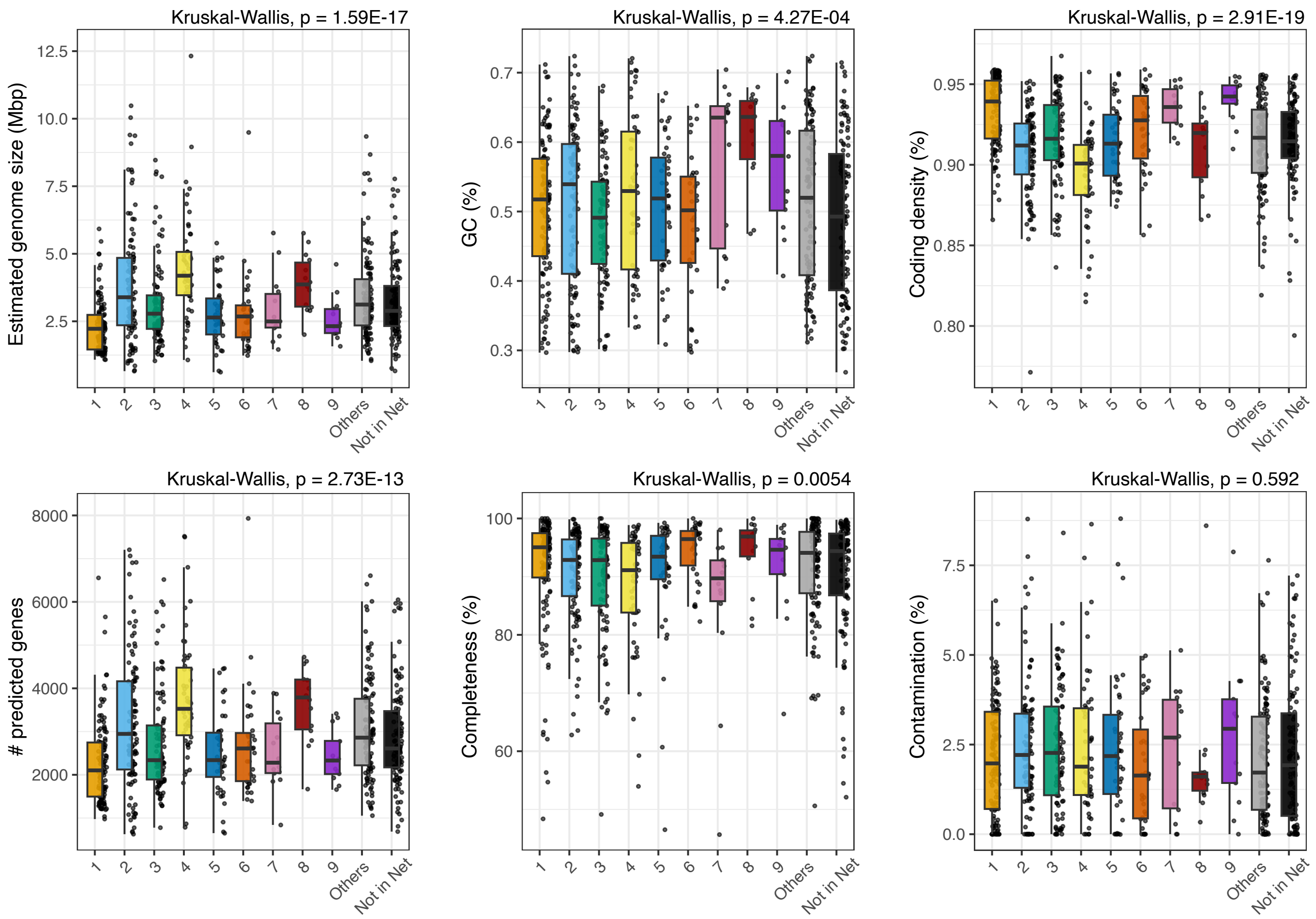


**Supplementary Figure 4. Genomic features of cohort members.** Boxplots showing the distribution of some genomic characteristics across cohorts, as well as species classified as "Others" and "Not in network". (A) Estimated genome size (Mbp), (B) GC content (%), (C) coding density (%), (D) number of predicted genes, (E) genome completeness (%), and (F) genome contamination (%). Points represent individual species. Boxplots show median and interquartile range, with whiskers extending to 1.5× IQR. Kruskal–Wallis test *P* values are shown above each panel.

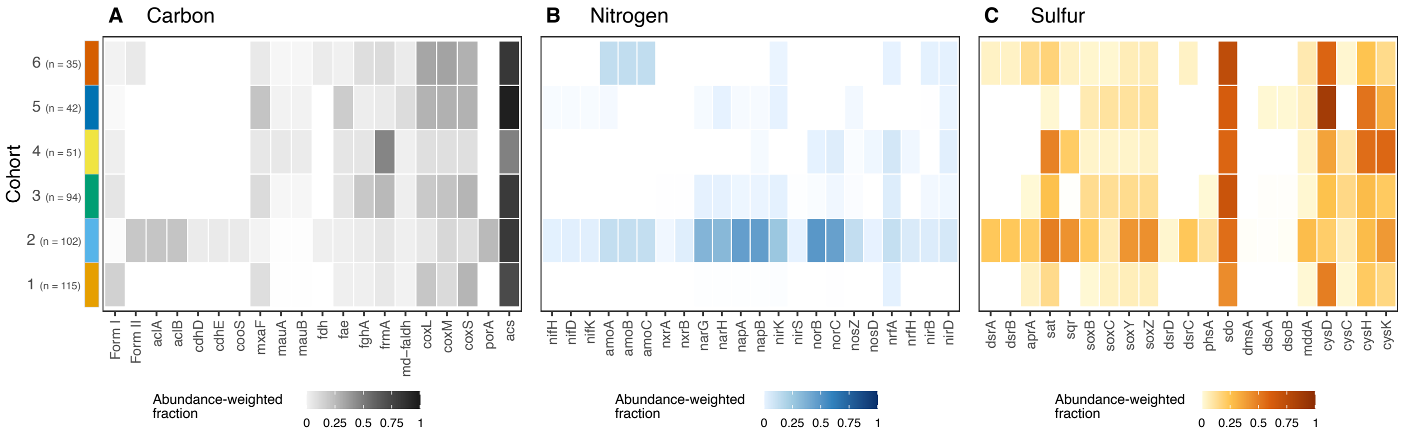


**Supplementary Figure 5. Relative abundance contribution to carbon, nitrogen, and sulfur genes across Baltic Sea microbial cohorts.** Heatmaps show the fraction of cohort-level mean relative abundance carried by genomes encoding metabolic genes involved in the (A) carbon, (B) nitrogen, and (C) sulfur biogeochemical cycles, based on METABOLIC annotations. Rows correspond to microbial cohorts, and columns represent functional genes for each of the main biogeochemical cycles. Gene descriptions are provided in Supplementary Table X. Color intensity (0–1) indicates the proportion of cohort abundance associated with genomes encoding the respective gene. Values are normalized within each cohort.

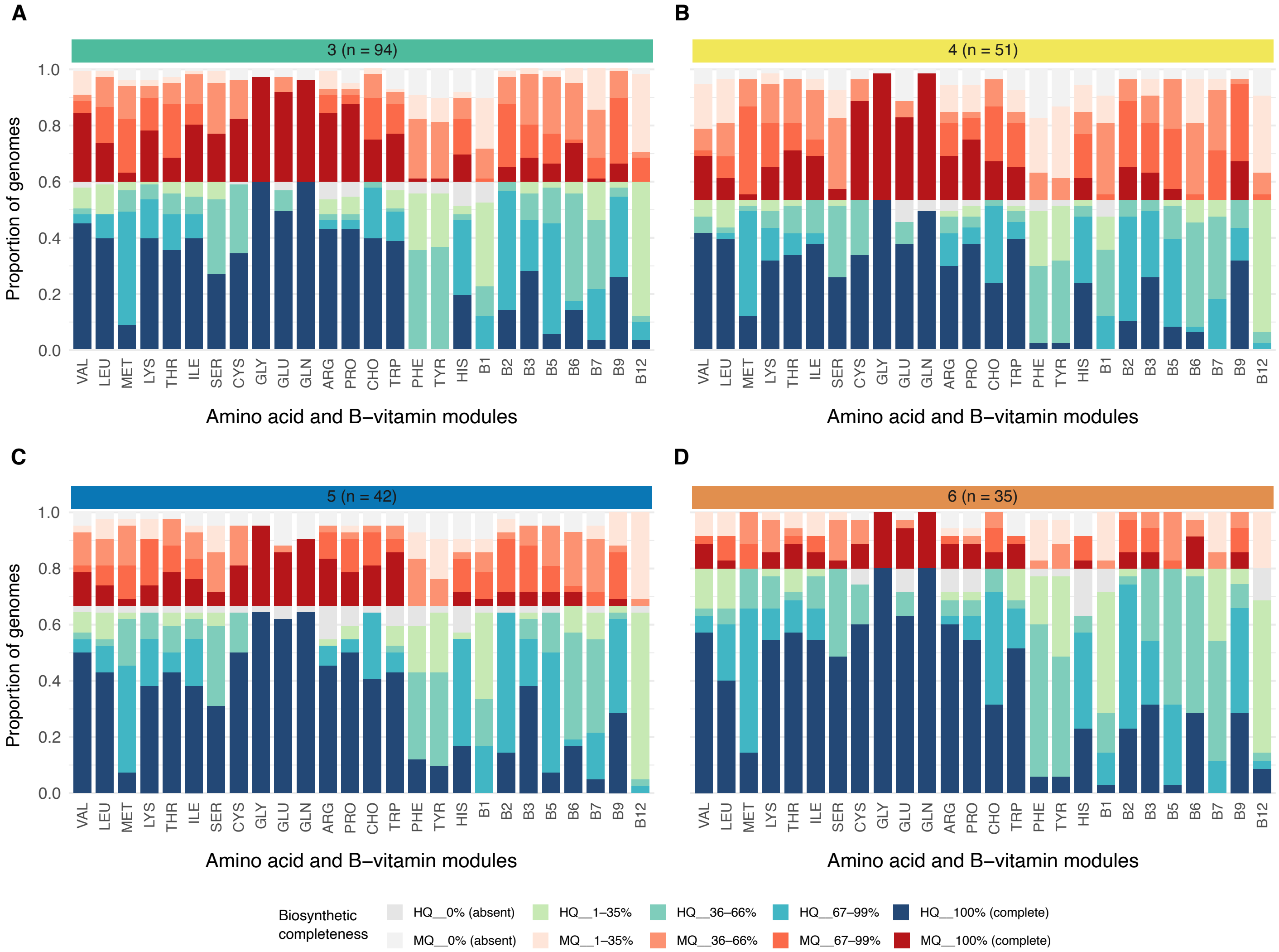


**Supplementary Figure 6. Amino acid and B-vitamin biosynthetic pathways across Baltic Sea microbial cohorts.** Custom-defined module completeness for eighteen amino acids and nine B-vitamins across members of (A) cohort 3, (B) cohort 4, (C) cohort 5, and (D) cohort 6. Species were grouped by genome quality into high-quality (HQ; >90% completeness and <5% contamination) and medium-quality (MQ; ≥50% completeness and <10% contamination) genomes. Stacked bar plots show the proportion of genomes across biosynthetic completeness categories for each pathway. Within each cohort, bars represent individual pathways, and the y-axis indicates the proportion of genomes assigned to each completeness group, further separated by genome quality (HQ at the bottom; MQ above). Biosynthetic completeness categories are defined as 0% (absent), 1–35%, 36–66%, 67–99%, and 100% (complete). Five color shades represent completeness levels for HQ genomes, and another five shades correspond to levels for MQ genomes. Numbers above each panel indicate the number of high-quality genomes relative to the total number of species within that cohort.

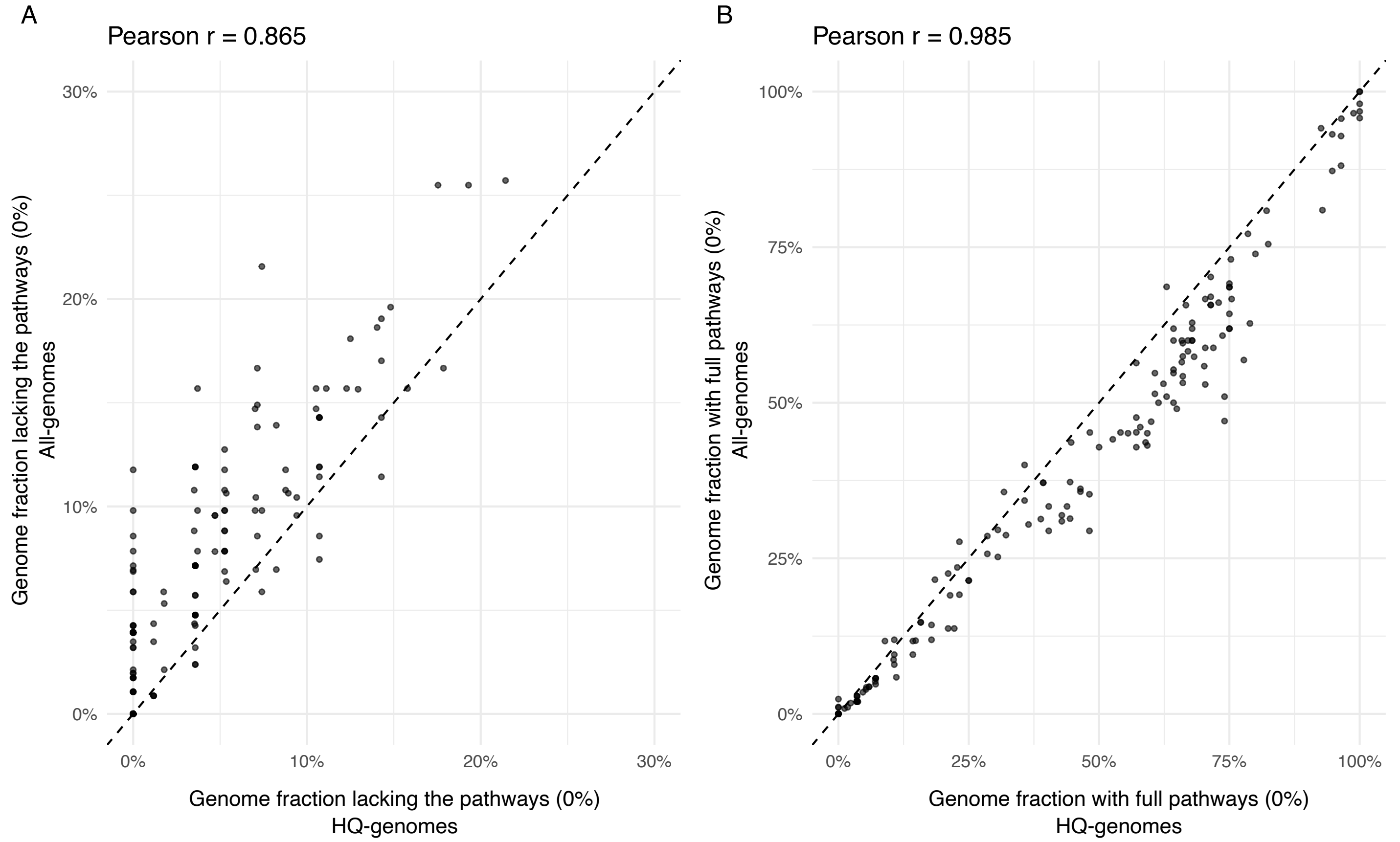


**Supplementary Figure 7. Comparison of pathway completeness estimates between high-quality genomes and the full genome dataset.** Scatter plots comparing pathway completeness using only high-quality genomes (>90% completeness, <5% contamination) against all genomes. Each point represents the fraction of genomes that (A) lack the pathway or (B) encode a complete pathway. Dashed lines indicate the 1:1 relationship. Pearson correlation coefficients are shown for each comparison.

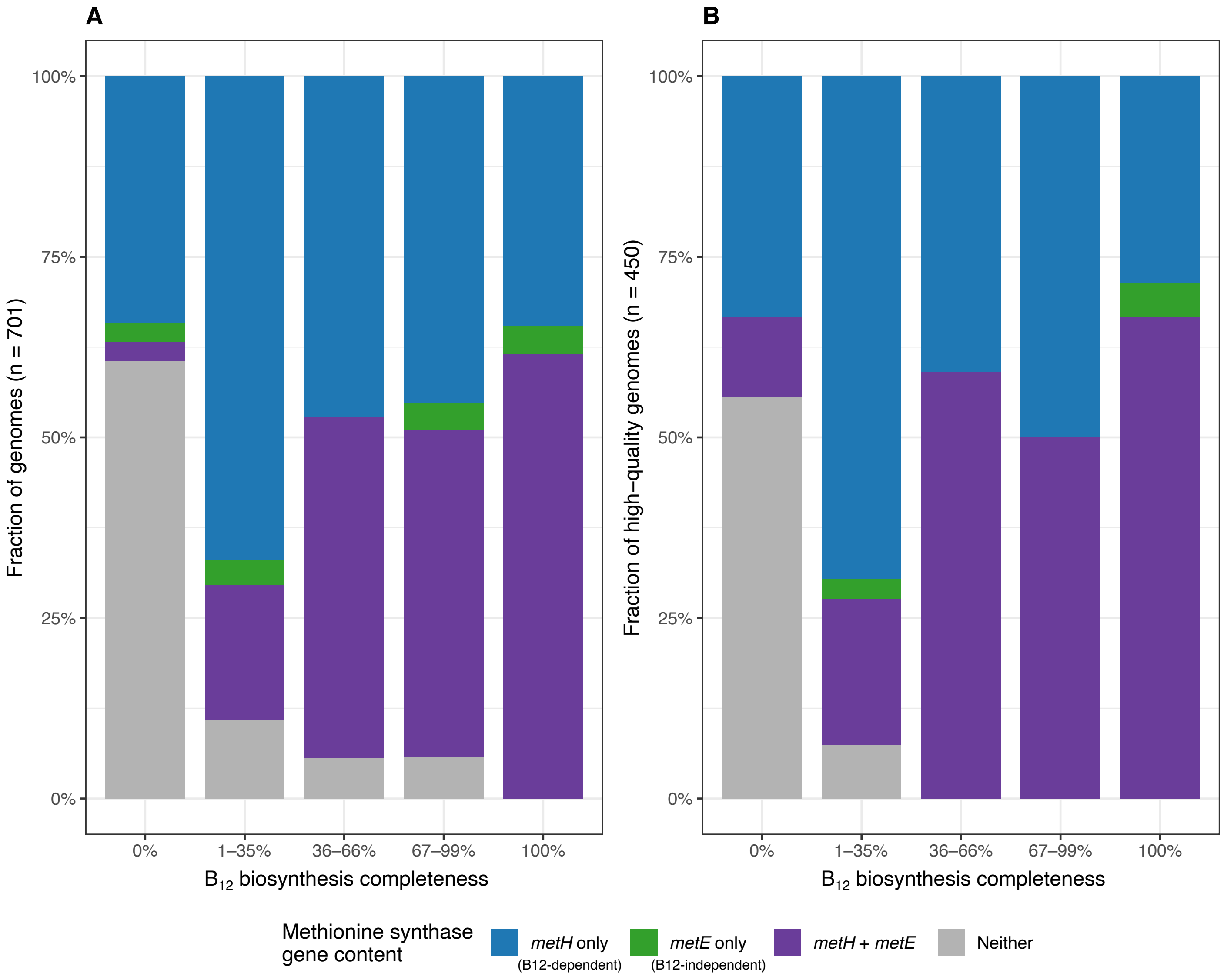


**Supplementary Figure 8. Distribution of methionine synthesis strategies across $B_{12}$ biosynthesis completeness categories.** (A) Stacked bar plot showing the fraction of genomes assigned to four methionine synthesis strategies across categories of best-route $B_{12}$ biosynthesis completeness: 0% (absent), 1–35%, 36–66%, 67–99%, and 100% (complete) for the full Baltic Sea MAG dataset (n = 701). Methionine strategies were defined based on the presence of the methionine synthase genes *metH* and *metE*: *metH* only, *metE* only, *metH* + *metE*, or "Neither". Best-route $B_{12}$ completeness was defined for each genome as the higher of the two custom $B_{12}$ biosynthesis module values, representing the aerobic or anaerobic route, each of which combines corrin ring biosynthesis with nucleotide loop assembly. (B) Same analysis restricted to high-quality genomes (n = 450; >90% completeness and <5% contamination).
